# *Caenorhabditis elegans* LET-413 Scribble is essential in the epidermis for growth, viability, and directional outgrowth of epithelial seam cells

**DOI:** 10.1101/2021.04.11.439327

**Authors:** Amalia Riga, Janine Cravo, Ruben Schmidt, Helena R. Pires, Victoria G. Castiglioni, Sander van den Heuvel, Mike Boxem

**Affiliations:** Division of Developmental Biology, Institute of Biodynamics and Biocomplexity, Department of Biology, Faculty of Science, Utrecht University, Utrecht, The Netherlands

**Keywords:** *C. elegans*, LET-413, Scribble, DLG-1, Discs large, cell polarity, migration, outgrowth, seam cells

## Abstract

The conserved adapter protein Scribble (Scrib) plays essential roles in a variety of cellular processes, including polarity establishment, proliferation, and directed cell migration. While the mechanisms through which Scrib promotes epithelial polarity are beginning to be unraveled, its roles in other cellular processes including cell migration remain enigmatic. In *C. elegans*, the Scrib ortholog LET-413 is essential for apical–basal polarization and junction formation in embryonic epithelia. However, whether LET-413 is required for postembryonic development or plays a role in migratory events is not known. Here, we use inducible protein degradation to investigate the functioning of LET-413 in larval epithelia. We find that LET-413 is essential in the epidermal epithelium for growth, viability, and junction maintenance. In addition, we identify a novel role for LET-413 in the polarized outgrowth of the epidermal seam cells. These stem cell-like epithelial cells extend anterior and posterior directed apical protrusions in each larval stage to reconnect to their neighbors. We show that the role of LET-413 in seam cell outgrowth is mediated at least in part by the junctional component DLG-1 discs large, which appears to restrict protrusive activity to the apical domain. Finally, we demonstrate that the Rho-family GTPases CED-10 Rac and CDC-42 can regulate seam cell outgrowth and may also function downstream of LET-413. Our data uncover multiple essential functions for LET-413 in larval development and shed new light on the regulation of polarized outgrowth of the seam cells.

## Introduction

Epithelial cells establish molecularly and functionally distinct apical, basal, and lateral membrane domains to function as selectively permeable barriers. Epithelial cell polarity is established through mutually antagonistic interactions between conserved groups of cortical polarity regulators, including the Par, Crumbs and Scribble modules (Benton and Johnston, 2003; Bilder et al., 2003; Laprise et al., 2006; Tanentzapf and Tepass, 2003). The Scribble polarity module, which consists of the proteins Scribble (Scrib), discs large (Dlg), and lethal giant larvae (Lgl), plays conserved roles in the establishment of basolateral identity and formation of cell junctions (Bossinger et al., 2001; Chalmers et al., 2005; Dow et al., 2003; Dow et al., 2007; Grifoni et al., 2007; Legouis et al., 2000; Mcmahon et al., 2001; Raman et al., 2016; Yamanaka et al., 2006). In addition, Scribble module proteins are involved in the regulation of cell proliferation and migration. In *Drosophila, scrib, dlg*, and *lgl* function as suppressors of neoplastic overgrowth of imaginal disks (Bilder et al., 2000; Gateff and Schneiderman, 1974; Mechler et al., 1985; Woods and Bryant, 1989). Many human tumors show altered Scrib protein levels or protein mislocalization, and in both *Drosophila* and mammalian tumor models, loss of Scrib increases the malignant and metastatic potential of oncogenic stimuli such as activation of Ras, Raf, Notch or Akt (Bonello and Peifer, 2019; Elsum et al., 2012; Santoni et al., 2020; Stephens et al., 2018).

The ability of Scrib to affect metastasis may be linked to its role as a regulator of cell migration. In *Drosophila, scrib* is required for migration of epithelial cells during dorsal closure (Bilder et al., 2003). In vertebrates, Scrib is involved in the migration of multiple cell types (Audebert et al., 2004; Dow et al., 2007; Michaelis et al., 2013; Nola et al., 2008; Osmani et al., 2006; Qin et al., 2005; Sun et al., 2017). How Scrib affects cell migration is not well understood, and may differ between cell types. In several epithelial cell types, Scrib is thought to regulate actin dynamics at the leading edge by promoting the recruitment or activation of the Rho-family GTPases Rac and Cdc42, or their effector proteins p21-activated kinases (PAKs) (Dow et al., 2007; Nola et al., 2008; Osmani et al., 2010). In other cell types, effects on cell migration do not appear to be mediated by small GTPases. In endothelial cells, Scrib regulates directed cell migration on fibronectin-coated surfaces by binding to and protecting surface integrin α5 from lysosomal degradation (Michaelis et al., 2013). In MDCK cells, Scrib affects cell migration through loss of E-cadherin-mediated cell adhesion (Qin et al., 2005). Finally, in dendritic cells and several cancer cell lines, Scrib was found to control cell migration downstream of the transmembrane semaphorin 4A (Sema4A) (Sun et al., 2017). In this context, cell migration appeared to be promoted by the downregulation of the activities of Scrib, Cdc42, and Rac1 (Sun et al., 2017). These examples illustrate the complexities of Scrib in cell migration.

The *C. elegans* genome encodes a single Scribble protein termed LET-413 that is essential for junction formation and epithelial polarization. In epithelia of embryos lacking *let-413* activity, junctional proteins fail to assemble into a continuous subapical belt and are found expanded through the lateral domain (Bossinger et al., 2001; Bossinger et al., 2004; KÖppen et al., 2001; Legouis et al., 2000; Mcmahon et al., 2001; Segbert et al., 2004). In addition, *let-413* embryonic epithelia show basolateral invasion of apical proteins including PAR-3, PAR-6, and the intermediate filament protein IFB-2 (Bossinger et al., 2004; Mcmahon et al., 2001). Ultimately, *let-413* embryos arrest due to a failure to elongate beyond the 1.5–2-fold stage. Investigating *let-413* in later stages of *C. elegans* development could provide further insights in the cellular pathways in which *let-413* participates, but the roles of *let-413* in larval development are not well characterized. Larval *let-413(RNAi)* causes sterility due to dysfunction of the spermathecal epithelium, where *let-413* was shown to be required for assembly of apical junctions and the maintenance of basolateral identify (Pilipiuk et al., 2009). However, no defects in growth rate or motility were reported. The role of *let-413* was also investigated in the intestine, using an intestine-specific CRISPR/Cas9 somatic mutant (Liu et al., 2018). Using this approach, LET-413 was shown to promote endocytic recycling. However, intestinal *let-413* CRISPR mutants continue larval development. Whether LET-413 is important in other larval tissues remains unclear. Moreover, it is not known if the role of Scrib in cell proliferation and migration is conserved in *C. elegans*.

Here, we use inducible protein degradation to investigate the roles of LET-413 in postembryonic epithelial tissues of *C. elegans*. Consistent with previous data, the presence of LET-413 in the intestine in larval stages was not essential for larval development. In contrast, we find that LET-413 is essential in the epidermis to support larval growth and viability, junctional integrity, and the directional outgrowth of the epithelial seam cells. The stem cell-like seam cells undergo an asymmetric division during each larval stage, followed by fusion of the differentiating anterior daughter cells with the surrounding epidermal syncytium. The remaining seam cells then form anterior–posterior directed protrusions of the apical domain to re-establish cell–cell contacts. To date, the mechanisms that mediate this dynamic shape change are poorly understood. Because of the essential role of LET-413 in junction maintenance, we investigated if the junctional component DLG-1 Discs large is also required for protrusion formation. We show that DLG-1 acts downstream of LET-413 and is essential for seam cell outgrowth, possibly by restricting protrusive activity to the apical domain. Nevertheless, seam cells depleted of DLG-1 show more protrusive activity than LET-413-depleted seam cells, indicating that regulating DLG-1 is not the only role of LET-413 in seam cell outgrowth. Finally, we investigated the contribution of the *C. elegans* Rac and Cdc42 GTPase family members to seam cell extension. We show that CED-10 Rac and CDC-42 can regulate seam cell outgrowth and may function downstream of LET-413. Together, our data indicate that the roles of Scrib in promoting protrusive cell shape changes are conserved in *C. elegans* and demonstrate essential roles for LET-413 Scrib in larval development and directed seam cell outgrowth.

## Results

### LET-413 is essential for larval growth and viability

To investigate the role of LET-413 in larval epithelia of *C. elegans*, we used the auxin inducible degradation (AID) system, which enables degradation of AID-degron-tagged proteins in a time- and tissue-specific manner (Nishimura et al., 2009; Zhang et al., 2015). We endogenously tagged LET-413 at the N-terminus shared by all predicted isoforms with the AID degron and a green fluorescent protein (GFP) (Fig. 1A). Before morphogenesis, we detected LET-413 ubiquitously at cell membranes (Fig. 1B). From the bean stage onward, LET-413 localized to the basolateral membrane domain and at intercellular junctions of epidermal and intestinal cells (Fig. 1B). In larval stages and adults, LET-413 was expressed in the intestine, epidermis, excretory canal, and the reproductive system (vulva, uterus, and spermatheca), where it also appeared to localize to the basolateral membranes and cell junctions (Figs. 1C and S1). These localization patterns are consistent with previous studies using antibody staining and transgene expression (Legouis et al., 2000; Legouis et al., 2003; Mcmahon et al., 2001; Pilipiuk et al., 2009).

**Figure 1.**
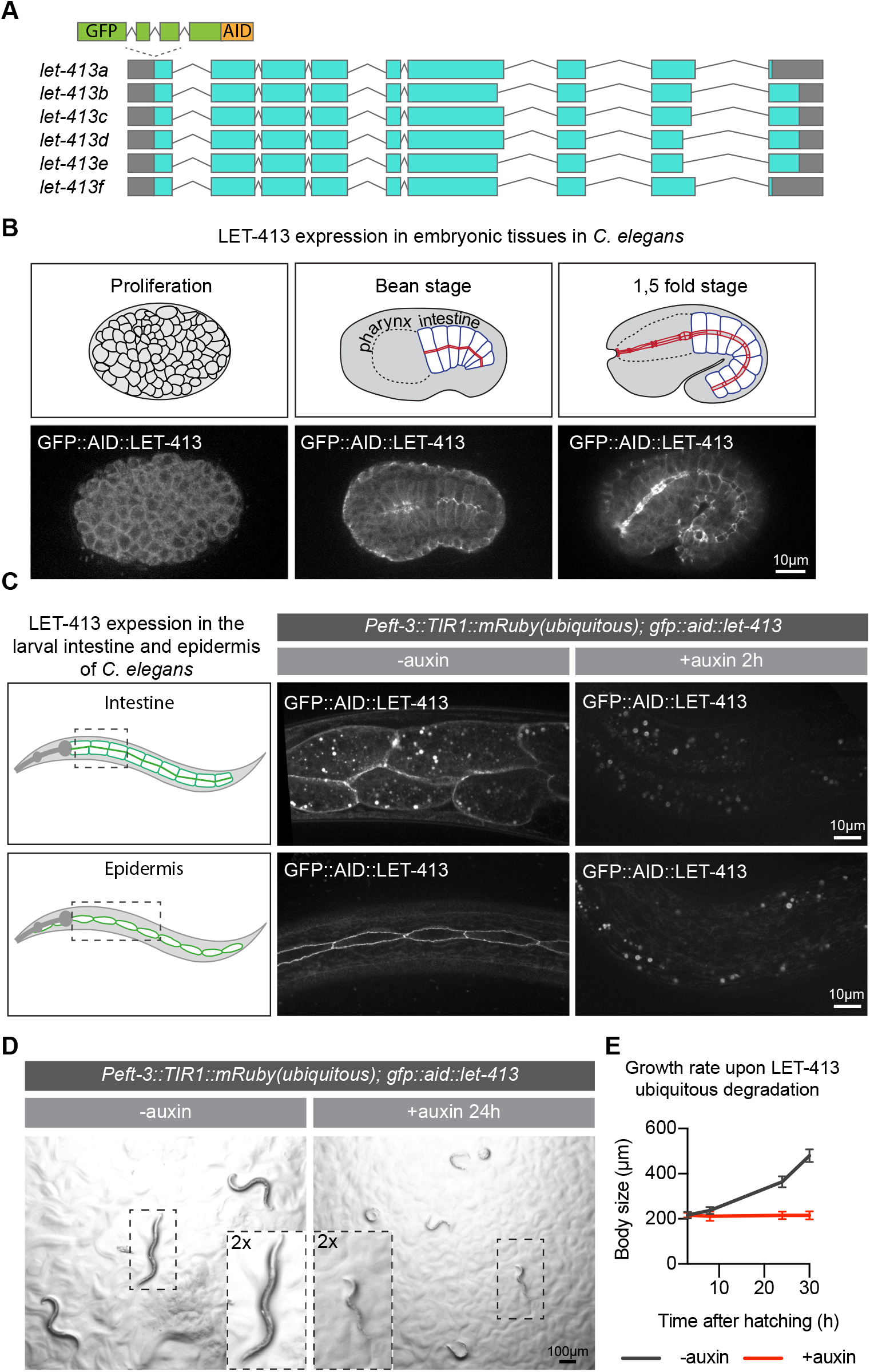
LET-413 is essential for larval development. (**A**) Schematic representation of predicted *let-413* splice variants and the insertion site of sequences encoding a green fluorescent protein (GFP) and the auxin-inducible degradation degron (AID). (**B**) Expression of LET-413 in embryonic development. (**C**) Expression of LET-413 in the intestine and epidermis of *Peft-3::TIR1::mRuby; GFP::AID::let-413* animals in the absence (-auxin) or presence (+auxin) of auxin for 2 h. Drawings are a schematic representation of the LET-413 expression pattern. Images are maximum intensity projections. In this and all other figures, 3 mM auxin is used (**D, E**) Growth of *Peft-3::TIR1::mRuby; GFP::AID::let-413* animals in the absence (-auxin) or presence (+auxin) of auxin. Images in D were acquired after 24 h of exposure to auxin. Growth curves in E show mean length ± SD upon continuous exposure to auxin. N = 8, 10, 12, and 9 for -auxin; and 10, 14, 15, and 13 for +auxin.

To determine whether LET-413 is essential for larval development, we degraded LET-413 using ubiquitously expressed TIR1 under control of the *eft-3* promoter, which is active in most or all tissues during larval development (Zhang et al., 2015). After 2 h of exposure of L1 larvae to auxin, LET-413 levels were depleted throughout the animal body. Quantifications of the GFP levels at epidermal cell junctions and the basolateral domain of intestinal cells revealed loss of GFP enrichment after auxin addition, confirming efficient degradation of LET-413 (Fig. 1C). Degradation of LET-413 from hatching onward resulted in a growth arrest and larval lethality (Fig. 1D, E). At 6 h of development, LET-413-depleted animals were already measurably smaller than control animals not treated with auxin (Fig. 1E). At 24 h after hatching, we observed not only a lack of growth, but also ∼80% larval lethality, as evidenced by lack of response to physical stimulation and lack of motility. These results show that LET-413 is essential for larval development.

### LET-413 is essential in the larval epidermis, but not the intestine

We next wanted to determine which larval tissue(s) contribute to the observed growth defect and lethality. We focused on two major larval epithelial tissues: the intestine and the epidermis. To deplete LET-413 specifically in these tissues, we made use of single-copy integrated lines expressing TIR1 from the tissue-specific promoters *Pelt-2* and *Pwrt-2,* which are active in the intestine and epidermis, respectively (Castiglioni et al., 2020). We examined intestinal morphology using an endogenous fusion of mCherry to the junctional protein DLG-1 Discs large but did not observe any defects in the characteristic junctional localization pattern, despite a lack of detectable LET-413 (Fig. 2A–C). Animals depleted of LET-413 in the intestine also grew normally (Fig. 2D). These data indicate that intestinal functioning does not critically depend on the continuous presence of LET-413. We next tested the requirement for LET-413 in the epidermis. Degradation of LET-413 occurred rapidly, with no GFP signal detected in the junctions of seam cells 2 h after the addition of auxin (Fig. 2E, G). In contrast to the intestine, degradation of LET-413 in the epidermis resulted in severe growth defects and larval lethality (Fig. 2D, F). Compared to ubiquitous degradation of LET-413, the growth defect was slightly less severe and more variable, while the larval lethality at 24 h of development was similar (Fig. 2H). These data show that LET-413 is essential for the functioning of the larval epidermis in *C. elegans*.

**Figure 2.**
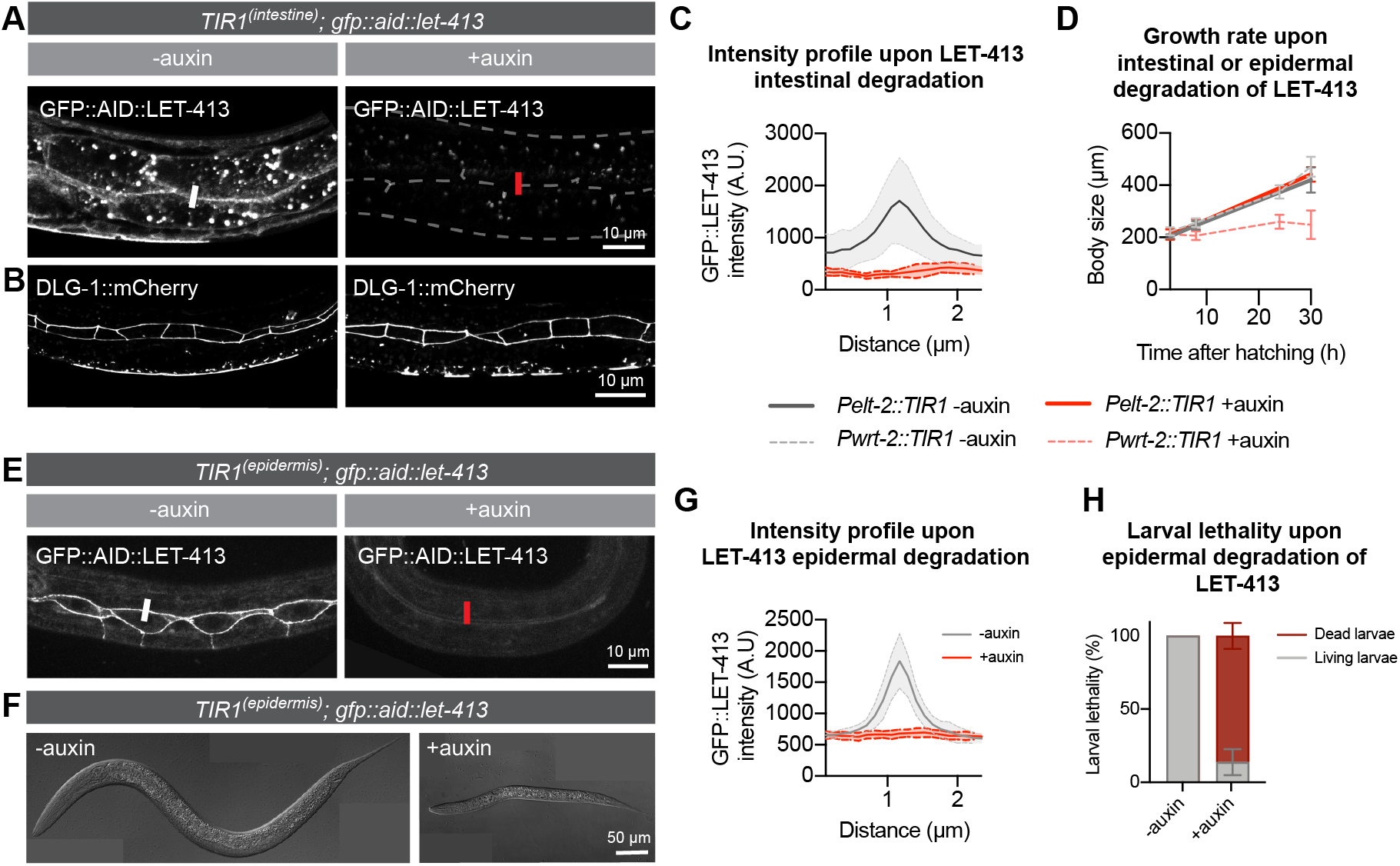
LET-413 is essential in the larval epidermis, but not the intestine. (**A**) Distribution of LET-413 in the intestines of *Pelt-2::TIR1::BFP; GFP::AID::let-413; dlg-1::mCherry* animals in the absence (-auxin) or presence (+auxin) of auxin for 36 h. Images are a single focal plane at the level of junctions between opposing intestinal cells. White and red bars are an example of sites used for fluorescence intensity quantification in C. (**B**) Distribution of DLG-1::mCherry in the same animals as in A. Images are maximum intensity projections of the intestinal cell junctions. (**C**) Intensity profiles of GFP::LET-413 fluorescence at junctions of opposing intestinal cells in *Pelt-2::TIR1::BFP; GFP::AID::let-413; dlg-1::mCherry* animals in the absence (-auxin) or presence (+auxin) of auxin for 30 h. Solid lines and shading represent mean ± SD. N=4 for both conditions. (**D**) Growth curves of *Pelt-2::TIR1::BFP; GFP::AID::let-413* and *Pwrt-2::TIR1::BFP; GFP::AID::let-413* animals in the absence (-auxin) or presence (+auxin) of auxin. Solid lines represent animals that express TIR1 in the intestine (*Pelt2::TIR1::BFP*) and dashed lines in the epidermis (*Pwrt2::TIR1::BFP*). Data show mean ± SD. N = 10, 13, and 10 animals for *Pelt2::TIR1* -auxin; 13, 13, and 10 for *Pelt2::TIR1* +auxin; 14, 14, 13 and 10 for *Pwrt2::TIR1* -auxin; and 14, 16, 18, and 11 for *Pwrt2::TIR1* +auxin. (**E**) Distribution of LET-413 in the epidermis of *Pwrt-2::TIR1::BFP; GFP::AID::let-413* animals in the absence (-auxin) or presence (+auxin) of auxin for 2 h. White and red bars are an example of sites used for fluorescence intensity quantification in G. (**F**) Example of growth of *Pwrt-2::TIR1::BFP; GFP::AID::let-413* animals in the absence (-auxin) or presence (+auxin) of auxin for 24 h. (**G**) Intensity profiles of GFP::LET-413 fluorescence at seam–hyp7 junctions in *Pwrt-2::TIR1::BFP; GFP::AID::let-413* animals in the absence (-auxin) or presence (+auxin) of auxin for 2 h. Solid lines and shading represent mean ± SD. N=4 animals for -auxin and 5 for +auxin. (**H**) Percentage of larval lethality upon degradation of epidermal LET-413 in *Pwrt-2::TIR1::BFP; GFP::AID::let-413* larvae grown in the presence of auxin from hatching. Data show mean ± SD.

### LET-413 is required for seam cell extension and reattachment

The epidermis of *C. elegans* larvae largely consists of the syncytial hypodermal cell hyp7 and two lateral rows of end-to-end attached epithelial seam cells embedded within hyp7 (Fig. 3A). Animals hatch with a complement of 10 seam cells (H0–H2, V1–V6 and T) (Fig. 3A), which undergo a reproducible pattern of asymmetric and symmetric divisions at specific times in development (Fig. 3B) (Sulston and Horvitz, 1977). In each larval stage, the V1–V4 and V6 seam cells undergo an asymmetric division, which generates an anterior daughter that differentiates and fuses with the hypodermis, while the posterior daughter retains the seam cell fate. In the second larval stage, the asymmetric division is preceded by a symmetric division that generates two seam cells. V5 follows a similar division pattern, except for the anterior daughter of the L2 division which becomes a neuroblast that generates a sensory structure termed the posterior deirid sensillium. Following the asymmetric divisions and fusions of the anterior daughters, the remaining seam cells extend anterior and posterior protrusions towards their neighbors and reattach, closing the gaps left by the fused cells.

**Figure 3.**
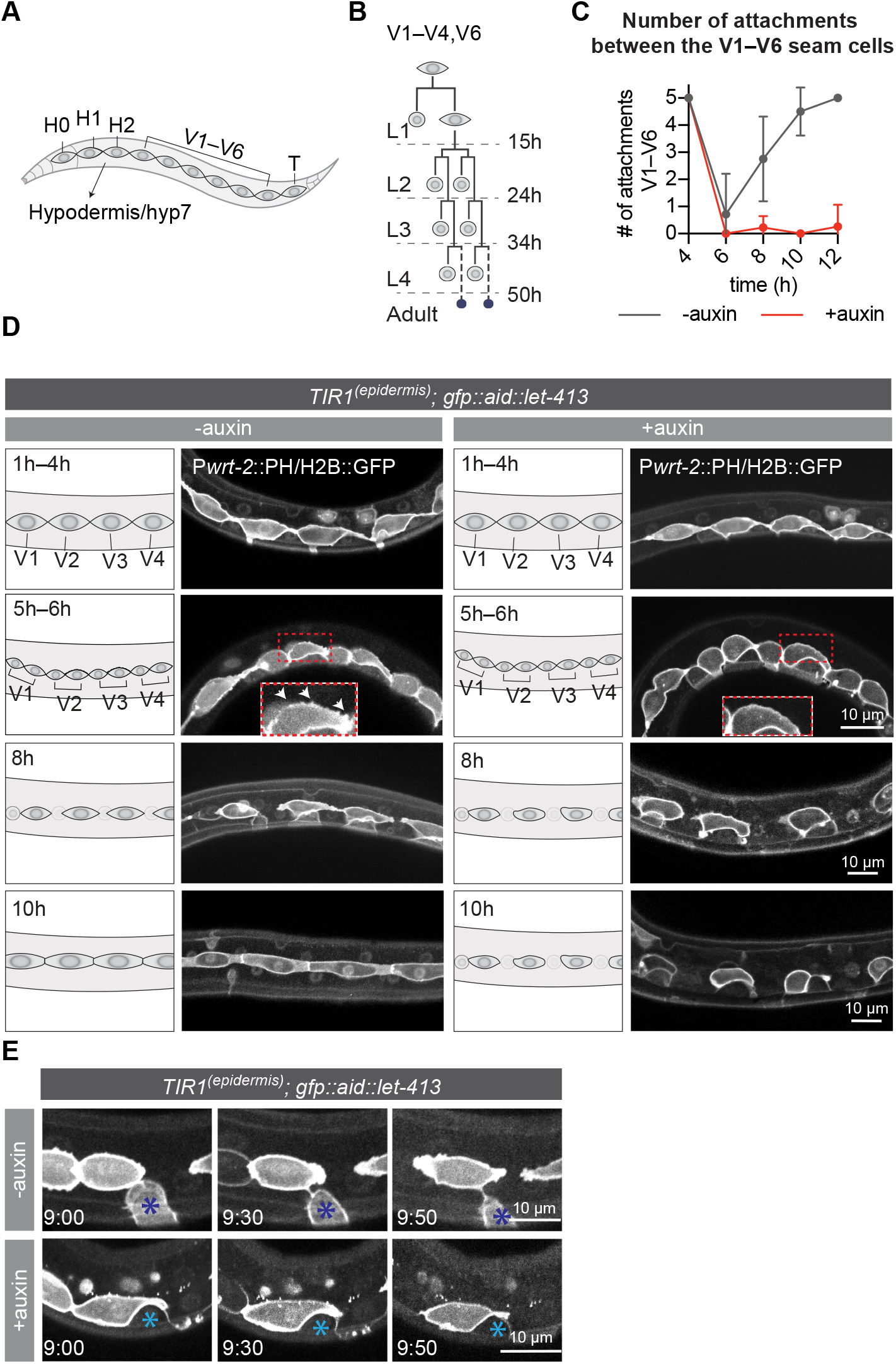
Loss of LET-413 leads to failure in seam cell outgrowth and reattachment after the asymmetric divisions. (**A**) Schematic representation of the *C. elegans* epidermis (lateral view). (**B**) Division pattern of the V1–V4 and V6 seam cells. Dotted lines demarcate the larval stages. Asymmetric divisions generate one differentiating daughter cell (round anterior cell) and one seam cell. During the L4 to adult molt, the remaining seam cells fuse and form a syncytium (small blue cells). (**C**) Number of attachments between the V1–V6 seam cells in LET-413-depleted (+auxin) or control animals (-auxin). Solid lines represent mean ± SD. N = 7 for -auxin time points, and 7, 9, 9, and 8 for +auxin time points. Genotype is *Pwrt-2::TIR1::BFP; GFP::AID::let-413; heIs63 [Pwrt-2::GFP::PH; Pwrt-2::GFP::H2B; Plin-48::mCherry]*. (**D**) Time series of L1 seam cell divisions and subsequent extension in LET-413-depleted (+auxin) or control animals (-auxin). Seam-specific GFP::H2B and GFP::PH mark DNA and cell membrane, respectively. Images are single apical focal planes. White arrows at 5–6 h time point indicate membrane protrusions. (**E**) P cell retraction in LET-413-depleted (+auxin) or control animals (-auxin). Images were taken at 9 h post hatching +0 min, +30 min, and +50 min. Blue asterisks indicate P cells. Seam-specific GFP::H2B and GFP::PH mark DNA and cell membrane, respectively.

To understand how loss of LET-413 affects the epidermal epithelium, we followed the seam cell division pattern in animals treated with auxin from hatching, using seam cell-specific GFP reporters that mark the cell membrane (*Pwrt-2::GFP::PH^PLC1d^*) and DNA (*Pwrt-2::GFP::H2B*) (Wildwater et al., 2011). Degradation of LET-413 did not affect the asymmetric seam cell divisions or the fusion of the anterior daughters with hyp7 (Fig. 3D). Following cell fusion, however, the remaining seam cells failed to extend protrusions towards their neighbors and remained isolated (Fig. 3C, D). In control animals, the seam cells showed signs of active membrane dynamics immediately following cell division, displaying small filopodia-like protrusions around the cell body (Fig. 3D, 5h–6h, white arrows). The apical domains of the seam cells then formed larger lamellipodium-like protrusive fronts directed towards the adjacent seam cells, and cells reattached around 10 h after hatching (Fig. 3C). Upon epidermal degradation of LET-413, the seam cells failed to form protrusions directed towards neighboring cells, and remained unattached (Fig. 3C, D). Depletion of LET-413 at later stages of development also resulted in isolated seam cells (Fig. S2). When depleting LET-413 just before the L2 divisions, we observed isolated clusters of >2 seam cells at a time where in control animals the posterior daughters had already reattached. This indicates that anterior seam cell daughters delay fusion or fail to fuse with hyp7, a defect not observed following L1 or L3 divisions. Presumably, the difference between larval stages is related to the unique division pattern of the seam cells in the L2 stage, in which the asymmetric divisions are preceded by a symmetric division that doubles the seam cell number. Nevertheless, LET-413 appears to be required for seam cell outgrowth throughout larval development.

Besides the seam cell extension defects, we also noticed defects in the retraction of the ventrolateral P cells (P1–P12). At hatching, 6 pairs of P cells cover the ventrolateral surface adjacent to V1 through V6 (Altun and Hall, 2009; Chisholm and Hsiao, 2012). Around the middle of the L1 stage, adjacent P cells separate from each other. Next, the P cells retract ventrally and interdigitate to form a single row of cells on the ventral side (Altun and Hall, 2009; Austin and Kenyon, 1994; Bone et al., 2016; Chisholm and Hsiao, 2012). In the presence of auxin, P cell pairs still became detached from their anterior and posterior neighbors but did not retract ventrally and appeared to stay in contact with the seam cells (Fig. 3E). This failure to retract may be accompanied by a change in cell fate or gene expression, as we noticed that LET-413 depletion results in reduced expression in P cells of the *Pwrt-2-*driven GFP::H2B and GFP::PH marker proteins (Fig. 3E). Finally, we tested whether LET-413 is required for migration of the Q neuroblasts, which are born within the lateral rows of seam cells but migrate anteriorly and posteriorly during L1 development (Rella et al., 2016). However, LET-413 is not expressed in the Q cell lineage, and we did not observe any defects in the migration of the Q cells or their descendants upon epidermal degradation of LET-413 (S3A, B).

Taken together, we conclude that LET-413 is required in the epidermis for outgrowth of the seam cells and retraction of the P cells, but not for migration of the Q cells.

### Loss of LET-413 causes junction impairment but does not affect the localization of apical PAR-6 or basolateral LGL-1

In embryonic epithelial tissues, LET-413 is important for junction assembly or integrity as well as for the basolateral exclusion of apical polarity determinants (Bossinger et al., 2001; Legouis et al., 2000; Mcmahon et al., 2001). To determine whether these roles are conserved in the larval epidermis, we examined the integrity of cell junctions and localization of apical–basal polarity markers upon depletion of LET-413. We first examined the distribution of the apical polarity regulator PAR-6 and the basolateral protein LGL-1. Under normal conditions, PAR-6 localizes to the apical surface of the seam cells, with enrichment at the cell junctions (Fig. 4A, B) (Castiglioni et al., 2020). Depletion of LET-413 did not affect the levels of PAR-6 at the apical and junctional domains (Fig. 4A, B), nor did it cause invasion of PAR-6 in the basolateral domain (Fig. 4C). We also did not detect apical invasion or reduced basolateral levels of LGL-1 (Fig. 4D, E). Thus, the depletion of LET-413 results in severe outgrowth defects, but does not appear to affect apical–basal polarity.

**Figure 4.**
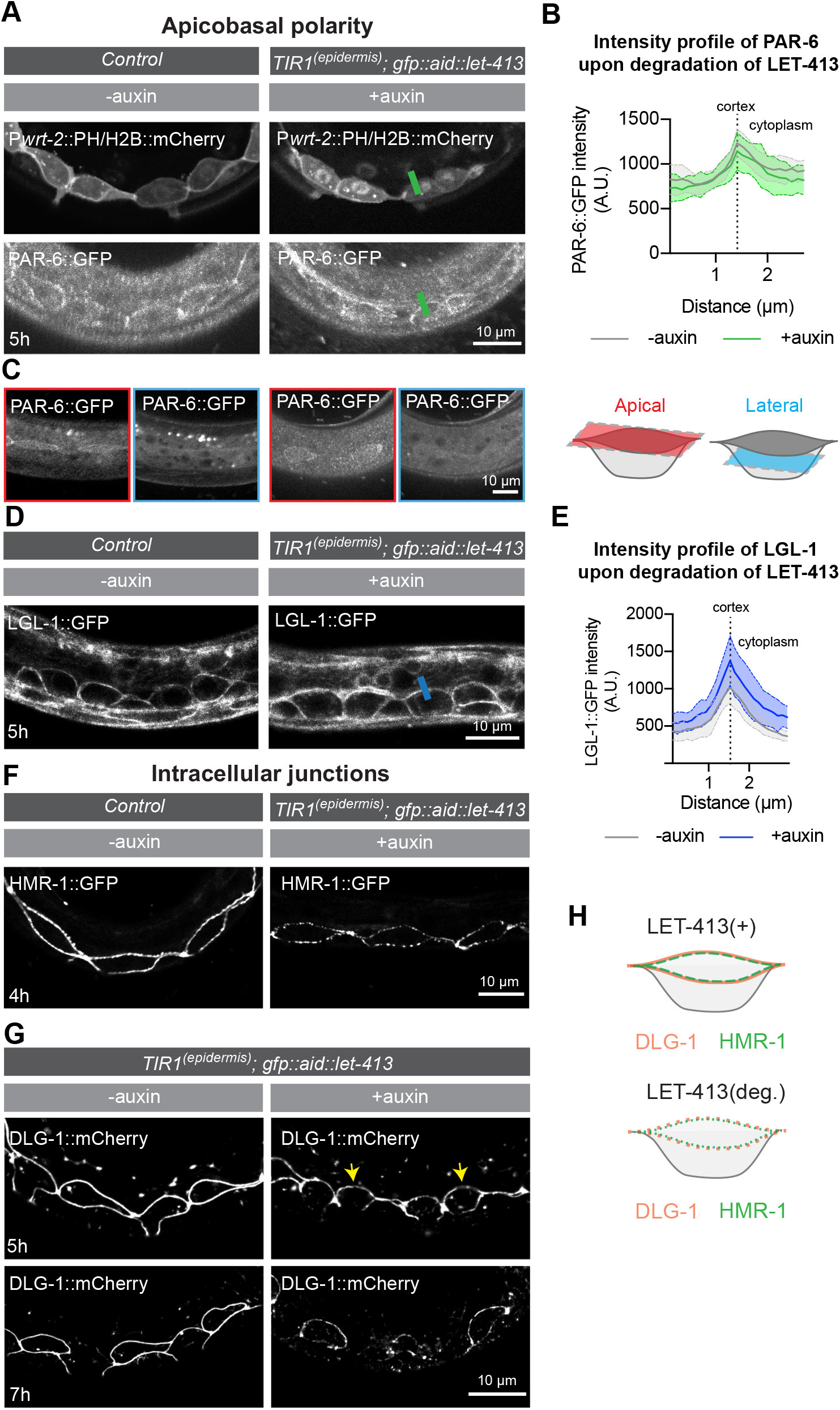
Loss of LET-413 causes junction impairment but does not affect the localization of the polarity regulators PAR-6 and LGL-1. (**A**) Distribution of PAR-6::GFP upon depletion of LET-413 at 5 h post hatching. As both PAR-6 and LET-413 are fused to GFP and their localization overlaps, the control animals do not carry the *Pwrt-2::TIR1::BFP* and *GFP::AID::let-413* gene fusions. (**B**) Quantification of the intensity profile of PAR-6::GFP in animals as in A. Graphs show mean apical GFP fluorescence intensity ± SD at the hyp7–seam cell junction (green bars in A). N = 5 animals for both conditions. (**C**) Distribution of PAR-6::GFP at the apical (red outlined panels) and lateral (blue outlined panels) domains of the seam cells. (**D, E**) Distribution and quantification of LGL-1::GFP in the epidermis of *lgl-1::GFP* animals without auxin and in *Pwrt-2::TIR1::BFP; GFP::AID::let-413; lgl-1::GFP* animals in the presence of auxin at 5 h post hatching. Images are maximum projections of the apical and junctional domain. Graphs show the mean junctional GFP fluorescence intensity ± SD at the hyp7–seam cell junction (blue bar in D). N = 5 animals. (**F**) Distribution of HMR-1 in the epidermis of *hmr-1::GFP* animals without auxin (-auxin) and *hmr-1::GFP; Pwrt-2::TIR1::BFP; GFP::AID::let-413* animals in the presence of auxin (+auxin) at 4 h post hatching. (**G**) Distribution of DLG-1::mCherry in the epidermis of *Pwrt-2::TIR1::BFP; GFP::AID::let-413; dlg-1::mCherry* animals without (-auxin) and in the presence of auxin (+auxin) at 5 and 7 h post hatching. Yellow arrows indicate punctate DLG-1::mCherry. (**H**). Schematic representation of HMR-1 and DLG-1 in the seam cells in the presence (+) or absence (-) of LET-413.

**Figure 5.**
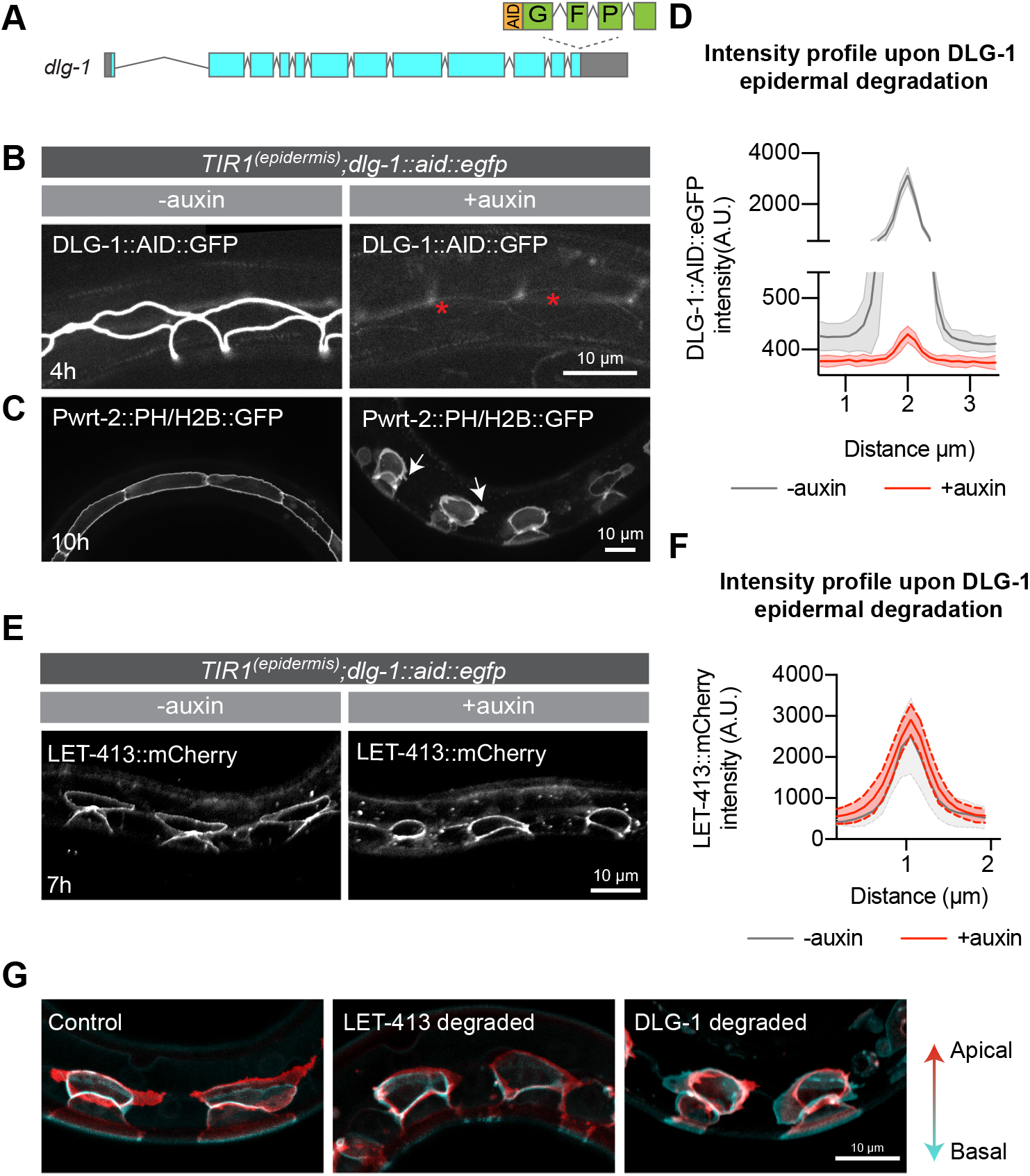
LET-413 acts upstream of DLG-1 in the regulation of the seam cell outgrowth. (**A**) Schematic representation of the insertion of GFP::AID-encoding sequences in the *dlg-1* locus. (**B, C**) Distribution of DLG-1 and PH::GFP and H2B::GFP markers, in control animals and DLG-1-depleted animals at indicated times post hatching. Genotypes are *Pwrt-2::TIR1::BFP; dlg-1::AID::GFP* for panel A, Pwrt-2::TIR1::BFP; *Pwrt2::ph::GFP Pwrt-2::H2B::GFP; dlg-1::AID::GFP* for panel B. Images are maximum projections of the apical domain. Asterisks indicate the position of DLG-1-depleted seam cells. Arrows indicate small apical protrusions, which were observed in >70% of cells in 10/10 animals examined. (**D**) Quantification of DLG-1 intensity across the hyp7–seam junction in animals depleted of DLG-1 as in panels B, C. Graphs show mean apical GFP fluorescence intensity ± SD. N = 8 animals for -auxin, and 4 animals for +auxin. (**E**) Distribution of LET-413 in control animals and DLG-1-depleted animals at indicated times post hatching. (**F**) Quantification of LET-413 intensity across the hyp7–seam junction in animals depleted of DLG-1 as in panels B, C. Graphs show mean apical GFP fluorescence intensity ± SD. N = 6 animals for -auxin, and 4 animals for +auxin. (**G**) Distribution of PH::GFP and H2B::GFP markers in control, LET-413-depleted and DLG-1-depleted seam cells at 8–10 h after hatching. Images are maximum projections of the entire cell body of the seam cells. The apical domains of the seam cells are indicated in red and the basal domains in cyan.

**Figure 6.**
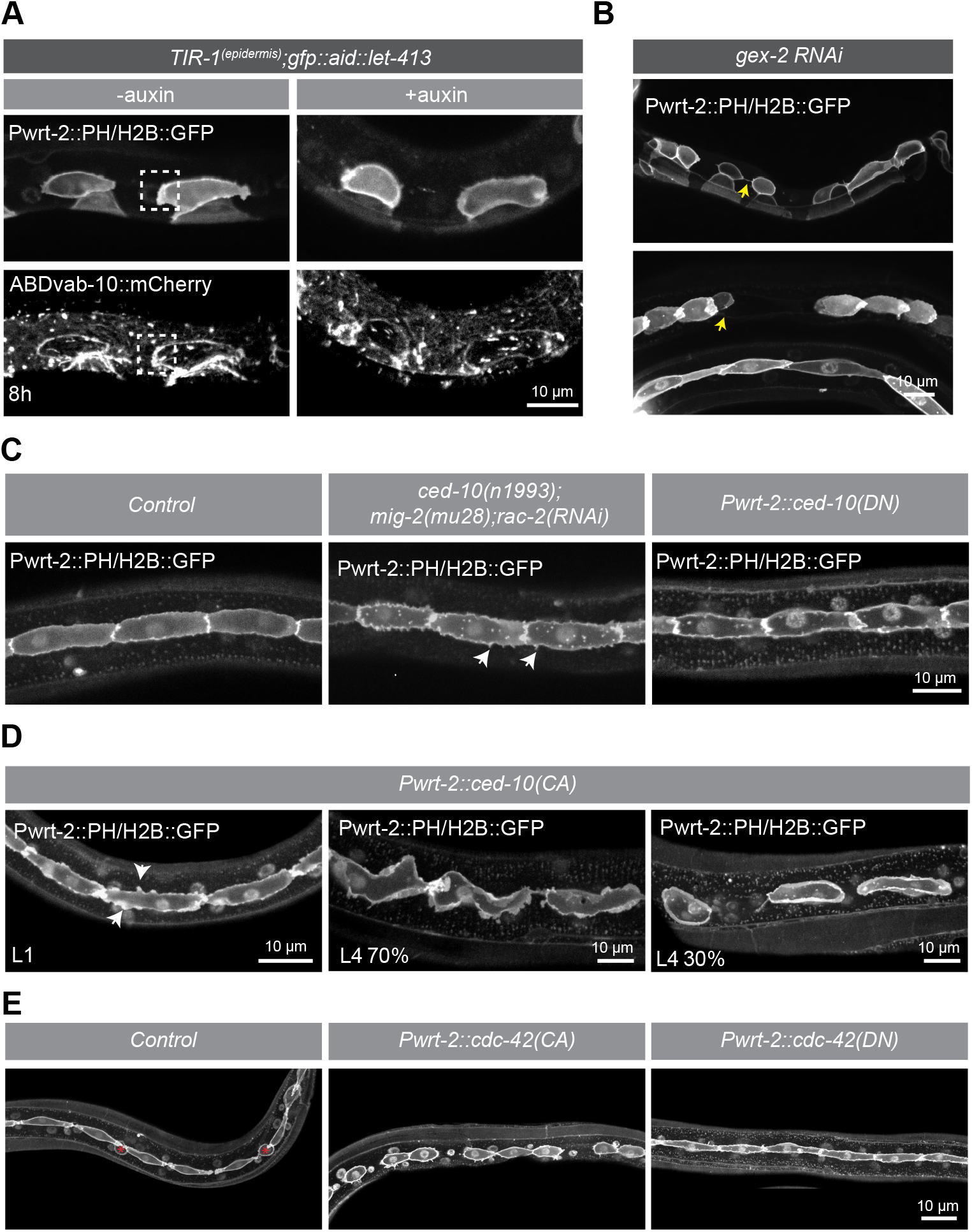
Involvement of Rac and Cdc42 GTPases in seam cell extension. (**A**) Distribution of the PH::mCherry and H2B::mCherry markers (van der Horst et al., 2019) and the VAB-10[ABD]::mCherry actin probe in the epidermis of *Pwrt-2::TIR1::BFP; GFP::AID::let-413* animals without (-auxin) and in the presence of auxin (+auxin) at 8 h post hatching. Images are single planes of the apical domain of the seam cells. White boxes indicate the enrichment of VAB-10[ABD] in the seam cell protrusions. (**B**) Distribution of the PH::GFP and H2B::GFP marker in *gex-2(RNAi)* animals. Yellow arrows point to defects in seam cell protrusions and seam cell shape. Defects were observed in 7/31 *gex-2(RNAi)* escaper animals examined. (**C**) Distribution of the PH::GFP and H2B::GFP markers in control, triple Rac GTPase depleted animals, and in animals expressing a dominant negative form of CED-10 Rac. White arrowheads indicate spike-shaped protrusions of the seam cells. (**D**) Distribution of the PH::GFP and H2B::GFP markers in animals expressing a constitutively active (CA) form of CED-10 in L1 and L4 larvae. White arrowheads point to representative cortex abnormalities in the seam cells. The percentages indicate the fraction of the imaged L4 animals showing the presented phenotype (N = 24 animals analyzed). (**E**) Distribution of the PH::GFP and H2B::GFP markers in control animals and animals expressing a constitutively active (CA) or dominant negative (DN) form of CDC-42 in L3 larvae. Asterisks indicate seam cells in the process of fusion with the hypodermis.

We next investigated the integrity of the *C. elegans* apical junctions (CeAJs) using a strain that expresses endogenously tagged HMR-1::GFP E-cadherin and DLG-1::mCherry Discs large, to mark the cadherin-catenin complex (CCC) and DLG-1/AJM-1 complex (DAC), respectively. DLG-1 forms a homogeneous junctional band around the seam cells, while HMR-1 shows a more punctate pattern (Fig. 4F, H). Epidermal depletion of LET-413 caused severe defects in the localization pattern of both HMR-1 and DLG-1. HMR-1 puncta became sparser, while DLG-1 was no longer localized in a continuous belt and instead localized to discontinuous short stretches and puncta (Fig. 4F–H). HMR-1 localization was already highly abnormal at 4 h post-hatching, well before the first asymmetric division of the seam cells (Fig. 4F). The first defects in DLG-1 localization were apparent at 5 h post-hatching and increased in severity over time (Fig. 4G). Together, our results demonstrate that the loss of LET-413 causes junction impairment but does not affect the apicobasal polarization of the seam cells.

### LET-413 acts upstream of DLG-1 in the regulation of seam cell outgrowth

During outgrowth, seam cells retain their overall apical–basal polarity and maintain the DLG-1/AJM-1 junctional complex. We therefore investigated if junctional defects contribute to the seam cell outgrowth defects of LET-413-depleted animals. To deplete DLG-1 specifically in the epidermis, we generated an endogenous C-terminal fusion of DLG-1 with AID::GFP (Fig. 5A). DLG-1 proved relatively resistant to auxin-mediated depletion. We therefore extended the auxin treatment by exposing newly hatched larvae to auxin for 24 h before initiating development. Using this approach, DLG-1 levels were reduced to ∼10% of non-auxin-treated controls (Fig. 5B, D). The seam cells in DLG-1 depleted animals showed severe outgrowth defects and did not extend towards the neighboring seam cells (Fig. 5C). This indicates that the seam cell outgrowth defects in LET-413 could be a consequence of the defects in DLG-1 organization. Alternatively, depletion of DLG-1 could affect the functioning of LET-413. However, depletion of DLG-1 did not affect the distribution or levels of LET-413 in the epidermis (Fig. 5E, F). Thus, in agreement with previous observations in the embryo and spermatheca (Bossinger et al., 2001; Bossinger et al., 2004; KÖppen et al., 2001; Legouis et al., 2000; Mcmahon et al., 2001; Pilipiuk et al., 2009; Segbert et al., 2004), LET-413 acts upstream of DLG-1 in the epidermis. Although DLG-1 depletion caused a failure of seam cell to extend and reattach, the phenotypes were less severe than those caused by LET-413 depletion. First, depletion of DLG-1 did not cause a growth arrest, though animals did appear to develop more slowly and became Dumpy. Second, small apical protrusions still formed (Fig. 5C, arrows). Finally, DLG-1 depletion caused an apparent expansion of the basolateral domains, as evidenced in maximum projections of Z-stacks of entire seam cells (Fig. 5G). These basolateral expansions occurred in an anterior–posterior direction, but whether this reflects active directional guidance or is due to physical constraints imposed by surrounding tissues is not clear. A possible explanation for the reduced severity of DLG-1 depletion compared to LET-413 depletion is that DLG-1 depletion was not fully complete. However, this does not explain the basolateral expansion caused by DLG-1 depletion. In addition, LET-413 depletion does not result in a complete loss of DLG-1 yet still causes a more complete block of seam cell outgrowth. Taken together, our data show that DLG-1 is essential for directed seam cell outgrowth and indicate that LET-413 regulates seam cell outgrowth in part by promoting the assembly or stability of DLG-1 at the CeAJ. However, the finding that depletion of LET-413 results in a more complete loss of protrusive activity than depletion of DLG-1 points to additional activities of LET-413 in promoting seam cell outgrowth.

### Actin dynamics and role of Rho-family small GTPases in seam cell extension

During seam cell extension, the cell membrane displays small filopodia-like protrusions, and a lamellipodium-like protrusive front forms at the anterior and posterior ends. It is likely therefore that reorganization of the actin cytoskeleton drives seam cell outgrowth. In migrating cells, members of the Rho family of small GTPases are essential to promote the cytoskeletal rearrangements needed for forward cell movement (Ridley, 2011; Schaks et al., 2019; Warner et al., 2019). One of the ways mammalian Scrib may promote cell migration is through regulating the activity or cortical localization of the Rac and Cdc42 members of the Rho family (Dow et al., 2007; Osmani et al., 2006). Moreover, inactivation of *C. elegans* CDC-42 by RNAi in larval stages was reported to result in defects in seam cell extension (Welchman et al., 2007). We therefore investigated if actin reorganization and regulation of Rac and Cdc42 may be important in the promotion of directed seam cell protrusions by LET-413.

We first examined the localization of actin during seam cell outgrowth, using an actin marker consisting of the actin-binding domain of VAB-10 fused to mCherry expressed in the hypodermis (*Plin-26::vab-10[ABD]::mCherry*) (Gally et al., 2009). We observed high levels of this actin probe in the protrusive fronts of extending seam cells (Fig. 6A, dashed boxes). In LET-413-depleted animals, in which seam cells do not develop a protrusive front, we did not observe polarized enrichment of actin (Fig. 6A). To determine if actin polymerization is required for protrusion formation, we inactivated the WAVE (<u>W</u>iskott-<u>A</u>ldrich syndrome protein family <u>ve</u>rprolin-homologous) complex component GEX-2 by RNAi feeding. *gex-2* function is required for Arp2/3-mediated branched actin nucleation in migrating cells (Sasidharan et al., 2018; Schaks et al., 2019). Because *gex-2(RNAi)* causes embryonic lethality (Soto etal., 2002), we placed adult animals on RNAi plates and examined rare escaper progeny. Animals also expressed VAB-10[ABD]::mCherry to discern RNAi affected from non-affected escapers. In escapers, we observed abnormal rounded seam cells and gaps between cells, indicative of a lack of seam cell extension (Fig. 6B, yellow arrows). Together, these observations indicate that seam cell extension is an actin-polymerization driven process.

Next, we investigated the role of the Rac and Cdc42 families of GTPases. The *C. elegans* genome encodes 3 Rac-related proteins, CED-10 Rac, MIG-2 RhoG, and RAC-2 Rac (Lundquist et al., 2001). Null alleles of *ced-10* are maternal effect embryonic lethal (Lundquist et al., 2001; Shakir et al., 2006; Soto et al., 2002). To investigate the effects of Rac inactivation, we combined the hypomorphic *ced-10(n1993)* allele with the predicted *mig-2(mu28)* null allele and *rac-2(RNAi)*. The combination of *ced-10(n1993)* with *mig-2(mu28)* was previously shown to result in a strong reduction of protrusive activity in intercalating dorsal epidermal cells in embryonic development (Walck-Shannon et al., 2015). However, seam cell outgrowth and reattachment still occurred in *ced-10(n1993); rac-2(RNAi); mig-2(mu28)* animals (Fig. 6C). We did notice an increase in filopodia-like protrusions (arrowheads in Fig. 6C), possibly due to a shift in the balance of activity between small GTPases. As an alternative approach to disrupt *ced-10* signaling, we expressed a dominant negative (DN) T17N mutant of CED-10 in the epidermis (Fig. 6C) (Bourne et al., 1991; Walck-Shannon et al., 2015). However, seam cells still developed protrusions, even when combining *ced-10(DN)* with *rac-2(RNAi)* and *mig-2(mu28)*. Thus, although we cannot exclude that some Rac activity remains, depletion of Rac function alone was not sufficient to prevent the formation of directed seam cell protrusions.

To investigate if activation of Rac signaling can promote protrusion formation, we also expressed a constitutively active (CA) Q61L mutant of CED-10 in the epidermis (Bourne et al., 1991; Walck-Shannon et al., 2015). To mark cells expressing CED-10, we used an expression cassette that also contains mCherry, separated from CED-10 by the T2A self-cleaving peptide. Expression of CED-10(CA) caused severe seam cell morphology defects, characterized by the appearance of excessive membrane ruffles or filopodia around the cell circumference (Fig. 6D). In older, L4 stage animals we also observed undirected protrusions of the seam cells (Fig. 6D, arrowheads). In most animals, anterior–posterior extension of the seam cells still formed and neighboring seam cells appeared to reattach. However, animals expressing high levels of mCherry showed rounded unattached seam cells (Fig. 6D, ∼30% of animals, N=24). These data support an important role for CED-10 Rac in regulating protrusion formation at the leading edge of extending seam cells.

We next examined the contribution of CDC-42, the single Cdc42 subfamily member present in *C. elegans*. As *C. elegans* CDC-42 is essential for embryonic development, we expressed dominant negative (DN) T17N and constitutively active (CA) Q61L mutants of CDC-42 in the epidermis (Bourne et al., 1991; Walck-Shannon et al., 2015). Expression plasmids also carried mTagBFP2, separated from CDC-42 by the F2A self-cleaving peptide. Expression of CDC-42(DN) did not result in defects in the extension of the seam cells (Fig. 6E). In contrast, expression of CDC-42(CA) resulted in rounded, detached seam cells (Fig. 6E and S4). The penetrance of the defects increased with age of the animals, ranging from 1 of 12 examined L1 animals to 30 of 34 L3 animals showing one or more gaps between the seam cells. Possibly this reflects accumulating CDC-42(CA) protein levels. Together, our data indicate potential roles for both CED-10 and CDC-42 in controlling the outgrowth of extending seam cells.

Finally, we wanted to investigate the effects of LET-413 depletion on CDC-42 GTPase activity. We expressed a CDC-42 fluorescent biosensor that specifically binds to the active GTP-bound form of CDC-42 in the epidermis (Kumfer et al., 2010). In control animals, we observed enrichment of the CDC-42 biosensor at the cortex of the seam cells, as well as in the anterior and posterior protrusions (Fig. S5B). A CDC-42::GFP fusion also localized to the protrusions (Fig. S5A). In LET-413-depleted animals, CDC-42::GFP remained enriched at the cell cortex but did not localize to protrusions (Fig. S6A, B). Although these results are consistent with LET-413 acting upstream to regulate CDC-42 activity, this cannot be concluded from these data as no protrusions are formed when LET-413 is depleted.

In summary, our data indicate an important role for the CED-10 Rac and CDC-42 GTPase proteins in reorganizing the actin cytoskeleton and driving anterior–posterior directed outgrowth of the seam cells. Although we cannot exclude an indirect role for LET-413 in this process, the observation that LET-413 depletion causes an almost complete lack of protrusive activity combined with prior findings that mammalian Scrib may regulate Rac and Cdc42, are consistent with LET-413 acting upstream of these small GTPases in seam cell extension.

## Discussion

The cortical polarity protein Scrib plays conserved roles in promoting basolateral identity and junction assembly in epithelial cells, and in regulating cell proliferation and migration. Studies of the single *C. elegans* Scrib homolog LET-413 identified essential roles for LET-413 in junction assembly and apical–basal polarization in the embryo and spermatheca, as well as a role in endocytic recycling in the intestine (Bossinger et al., 2001; Bossinger et al., 2004; KÖppen et al., 2001; Legouis et al., 2000; Liu et al., 2018; Mcmahon et al., 2001; Pilipiuk et al., 2009; Segbert et al., 2004). However, no essential roles in larval development or cell migration have been reported. Here, we used auxin-inducible protein degradation to bypass embryonic requirements and inactivate LET-413 in larval tissues. Using this approach, we find that expression of LET-413 in the larval epidermis is essential for growth and viability, and to promote directed outgrowth of the seam cells.

### LET-413 in growth and viability

The ubiquitous depletion of LET-413 from hatching onward caused a near complete lack of animal growth and resulted in larval lethality. Epidermal-specific depletion of LET-413 resulted in similar levels of larval lethality and a growth defect that was only slightly less severe. In contrast, we observed no growth defects or lethality when we depleted LET-413 in the intestine, consistent with previous observations of an intestine-specific CRISPR/Cas9 mutant of *let-413* (Liu et al., 2018). Thus, the growth and viability defects appear to be largely due to an essential requirement for LET-413 in the epidermis, while LET-413 is not required in the intestine for larval development or viability. The more severe growth defects observed upon ubiquitous LET-413 depletion may reflect a minor contribution of additional tissues. Alternatively, it remains possible that TIR1 driven by the ubiquitous *eft-3* promoter results in more effective epidermal LET-413 depletion than TIR1 driven by the epidermal *wrt-2* promoter, even though we did not observe a difference in depletion by microscopy.

The reason for the growth arrest following epidermal LET-413 degradation will require further investigation. It is not likely to be connected to the seam cell outgrowth defects, as the growth arrest is visible when the first seam cell divisions have not yet taken place. We recently found that epidermal depletion of the apical polarity regulators PAR-6 or PKC-3 similarly results in a larval growth arrest (Castiglioni et al., 2020). Thus, there may be a general requirement for apical–basal polarity regulators in the epidermis to support growth. Whether the growth defects are due to a loss of epidermal polarity or represent different functions of these polarity regulators remains to be determined.

### LET-413 in seam cell outgrowth

In LET-413-depleted animals, the posterior daughters of asymmetric seam cell divisions do not extend protrusions towards their neighboring cells, and consequently fail to reattach to each other. The process of directed seam cell extension has not been extensively studied. To our knowledge, the only genes specifically implicated in this process to date are *cdc-42* and *nhr-25* (Silhseam cells are somewhat similar to intercalatingÁnkovÁ et al., 2005; Welchman et al., 2007). CDC-42 may function downstream of LET-413 and is discussed further below. NHR-25 is a nuclear receptor family transcription factor whose inactivation causes a similar seam cell extension defect as LET-413 depletion (SilhÁnkovÁ et al., 2005). However, transcriptional targets through which NHR-25 controls seam cell outgrowth have not been identified, and *nhr-25* has numerous roles in *C. elegans* development in addition to seam cell outgrowth (Brooks et al., 2003; Mullaney et al., 2010; Shao et al., 2013; Ward et al., 2013). Nevertheless, it may be interesting to investigate if loss of *nhr-25* affects the transcription of *let-413*.

Because of the strong growth defects caused by LET-413 depletion, which are already apparent at the time of the first seam cell division, we considered whether the outgrowth defects are a secondary consequence of the growth arrest. However, we do not think that this is the case due to the highly specific nature of the seam cell phenotype. Division, fusion, and cell outgrowth all occur within a short time span, with outgrowth overlapping in time with the fusion process. Yet only outgrowth is affected by LET-413 depletion, and division and fusion take place as normal. Moreover, we did not observe a growth arrest in DLG-1-depleted animals, in which seam cells similarly fail to extend and reattach.

In a previous study using seam-specific RNAi, inactivation of *let-413* or the junction components *ajm-1* or *dlg-1* was postulated to cause a loss of seam cell fate and inappropriate fusion with the surrounding hypodermis, based on loss of an AJM-1::mCherry marker (Brabin et al., 2011). In our experiments with a PH::GFP membrane marker, we never observed fusion of posterior seam cells upon depletion of LET-413 or DLG-1. Nevertheless, depletion of LET-413 did not result in a complete loss of cell junctions nor in a complete loss of DLG-1 junction localization. It remains possible therefore that a stronger loss of junctional components could result in inappropriate fusion with the hypodermis.

LET-413 depletion also did not appear to affect the differentiation of the anterior daughter cells, as these fused with the hypodermis with normal kinetics. The only exception we observed followed the L2 seam cell divisions, with depletion of LET-413 starting in late L1 larvae. The division pattern in the L2 stage differs from the L1, L3, and L4 stages in that the asymmetric cell divisions are preceded by a symmetric division that increases the seam cell number. Thus, following the asymmetric, second division, four seam cells are generated, of which two will fuse with the hypodermis. At the time when fusion and reattachment of remaining seam cells were completed in control animals, the anterior seam cell daughters in LET-413 depleted animals had not fused. Possibly, LET-413 depletion affects aspects of seam differentiation that are particularly essential during the alternate L2 stage division pattern. Finally, the P cells in LET-413 depleted animals showed a reduced expression of fluorescent marker proteins driven by the *wrt-2* promoter. This may indicate a change in cell fate but could also be a secondary consequence of the failure in ventral retraction. Taken together, the loss of LET-413 may contribute to epidermal cell fate specification, but the primary defects we observe are in cell outgrowth and junctional integrity.

### DLG-1 and cell junctions in seam cell outgrowth

The loss of LET-413 in the epidermis led to impaired cell junctions, evidenced by the highly fragmented appearance of the junction components DLG-1 Discs large and HMR-1 E-cadherin. Similar junctional defects were demonstrated in embryonic epidermal cells and the spermatheca upon *let-413* inactivation (Bossinger et al., 2001; Bossinger et al., 2004; KÖppen et al., 2001; Legouis et al., 2000; Mcmahon et al., 2001; Pilipiuk et al., 2009; Segbert et al., 2004). We did not observe junctional defects in the intestine, indicating that junction maintenance in this tissue does not require the continued expression of LET-413. In a previous study, intestine-specific CRIPSPR/Cas9-mediated knockout of *let-413* was reported to cause lateral displacement of HMP-1 α-catenin (Liu et al., 2018). However, the *vha-6* promoter used to express Cas9 in this study is active during embryonic intestinal development, and the junctional defects may reflect a requirement in intestinal development. Our data show that LET-413 acts upstream of DLG-1 in junction maintenance in the seam epithelium, again in accordance with observations in the embryo and spermatheca (Bossinger et al., 2001; Bossinger et al., 2004; KÖppen et al., 2001; Legouis et al., 2000; Mcmahon et al., 2001; Pilipiuk et al., 2009; Segbert et al., 2004). In all *C. elegans* tissues examined to date, LET-413 acts upstream of DLG-1. This is different from *Drosophila*, where the localization hierarchy between the two proteins differs between tissues. In the adult *Drosophila* midgut epithelium, Scrib is required for the localization of Dlg (Chen et al., 2018), while in the follicle epithelium, Dlg acts upstream of Scrib (Khoury and Bilder, 2020).

One potential difference between our study and previous studies of *let-413* is the effect of LET-413 loss on apical–basal polarity. In embryonic epithelia, inactivation of *let-413* results in the basolateral invasion of apical proteins including PAR-3, PAR-6, and IFB-2 (Bossinger et al., 2004; Mcmahon et al., 2001). In the spermatheca, *let-413(RNAi)* also resulted in a lack of apical PAR-3. In contrast, we did not observe relocalization of the apical polarity protein PAR-6 or the basolateral polarity regulator LGL-1 upon LET-413 depletion. One explanation for this difference is that we investigate different tissues, and the epidermis may be less reliant on LET-413 for apical–basal polarization. We can also not rule out minor changes in localization not detectible with the markers and microscopy approaches used. The most likely explanation, however, is that the difference is caused by the timing of inactivation. While previous studies inactivated *let-413* from the start of embryonic development or prior to development of the spermatheca, we depleted LET-413 only after epidermal tissues are established. While the seam cells do continue to divide, established tissues may be less reliant on LET-413 expression for maintaining apical–basal domain identity.

Interestingly, directed outgrowth of the seam cells appears to depend on the DLG-1/AJM-1 junctional complex, as depletion of DLG-1 resulted in a lack of seam cell outgrowth. What could the role of cell junctions in this process be? Cell–cell junctions are essential in transducing mechanical forces generated by actomyosin contractions into coordinated morphogenetic changes during cell–cell intercalation and collective cell migration (Pannekoek et al., 2019; Pinheiro and BellaÏche, 2018; rauzi, 2020; Walck-Shannon and Hardin, 2014). The seam cells are somewhat similar to intercalating cells and cells undergoing collective migration in that they maintain their overall apical–basal polarity and cell junctions during outgrowth. However, force transduction is generally mediated by cadherin-based adherens junctions that, in contrast to the DLG-1/AJM-1 complex, directly connect to the actin cytoskeleton. A more likely possibility is that the DLG-1/AJM-1 junctional complex is involved in spatially restricting the protrusive activity to the apical domain. In support of this, upon loss of DLG-1 the basolateral sides of the cells expanded in an anterior–posterior direction. Potential mechanisms for such spatial restriction include functioning as a hub for signaling components, or even inappropriate paracellular passage of extracellular factors involved in protrusion formation like guidance cues. Finally, we cannot exclude that DLG-1 has additional roles not related to cell junctions.

### Actin and Rho GTPase family members in seam cell outgrowth

The almost complete lack of protrusive activity in LET-413-depleted animals indicated that LET-413 has additional functions in seam cell outgrowth besides the maintenance of the DLG-1 junctional complex. It is attractive to consider regulation of the actin cytoskeleton via the Rho-family small GTPases Rac and Cdc42 as an additional function of LET-413. For one, Rho-family GTPases play critical roles in coordinating the cytoskeletal rearrangements that drive cellular protrusions (Ridley, 2011; Schaks et al., 2019; Warner et al., 2019). *C. elegans* Rho-family GTPases have already been shown to be involved in many migratory events, such as long-range migration of the Q neuroblasts and the gonadal distal tip cells (Lundquist et al., 2001; Reddien and Horvitz, 2000; Rella et al., 2016), morphogenetic changes in embryonic epidermal cells during dorsal intercalation and ventral closure (Ouellette et al., 2016; Patel et al., 2008; Walck-Shannon et al., 2015; Walck-Shannon et al., 2016; Wallace et al., 2018; Zilberman et al., 2017), and growth cone migration in axonal pathfinding (Dyer et al., 2010; Lundquist et al., 2001; Shakir et al., 2006). In addition, RNAi-mediated inactivation of *cdc-42* was shown to prevent seam cell outgrowth (Welchman et al., 2007). Finally, mammalian Scrib is postulated to control cell migration in part through regulation of Rac and Cdc42 (Dow et al., 2007; Osmani et al., 2010).

Our data build upon these observations and support an important role for the regulation of actin dynamics by CED-10 Rac and CDC-42 in seam cell extension. Overexpression of constitutively active variants of these proteins resulted in increased and undirected protrusive activity as well as seam outgrowth failures, indicating that the spatial activity of these proteins needs to be carefully regulated for proper seam cell outgrowth and reattachment. Our data are consistent with LET-413 acting upstream of Rho-family GTPases. LET-413 may mediate the localization or activity of Rho GTPases themselves, of upstream regulators such as GTPase activating proteins or guanine nucleotide exchange factors, or of downstream effector proteins like the PAKs. However, given the many functions and interaction partners of Scrib proteins, LET-413 may also affect cytoskeletal dynamics quite indirectly, *e.g.,* by promoting the production, secretion, or response to migratory cues.

We were not able to block formation of seam cell protrusions through mutation, RNAi, or expression of dominant negative variants of Rac family members and *cdc-42*. We cannot be fully certain therefore that the activity of these proteins is essential for seam cell extension. A likely interpretation is that this is due to redundant activities. *C. elegans* expresses three Rac-family members, and in other migratory cell types redundancies have been shown between Rac family members themselves, as well as between Rac proteins and CDC-42 (Cao et al., 2021; Dyer et al., 2010; Lundquist et al., 2001; Peters et al., 2013; Walck-Shannon et al., 2015). Our inactivation experiments may therefore simply not have resulted in sufficient deregulation of Rac and Cdc42 signaling to disrupt seam cell extension. In support of this interpretation, expression of constitutively active CED-10 Rac and CDC-42 was able to block seam cell extension, indicating that sufficient disruption of GTPase signaling can block outgrowth. A further complicating factor in studying the role of small GTPases in seam cell extension is that they play numerous essential roles in *C. elegans* development. Potential future experiments in this direction would require the development of a more comprehensive toolkit that can inactivate any desired combination of Rho-GTPases with very high temporal precision.

In summary, our data show that LET-413 likely affects seam cell outgrowth through multiple pathways, including maintenance of the DLG-1 junctional complex and regulation of actin cytoskeleton dynamics. An important future direction will be to uncover how LET-413 regulates cytoskeletal dynamics.

## Materials and methods

### *C. elegans* strains and culture conditions

*C. elegans* strains were cultured under standard conditions (Brenner, 1974). Only hermaphrodites were used, and all experiments were performed with animals grown at 20 °C on Nematode Growth Medium (NGM) agar plates. Table 1 contains a list of all the strains used.

**Table 1 –.**
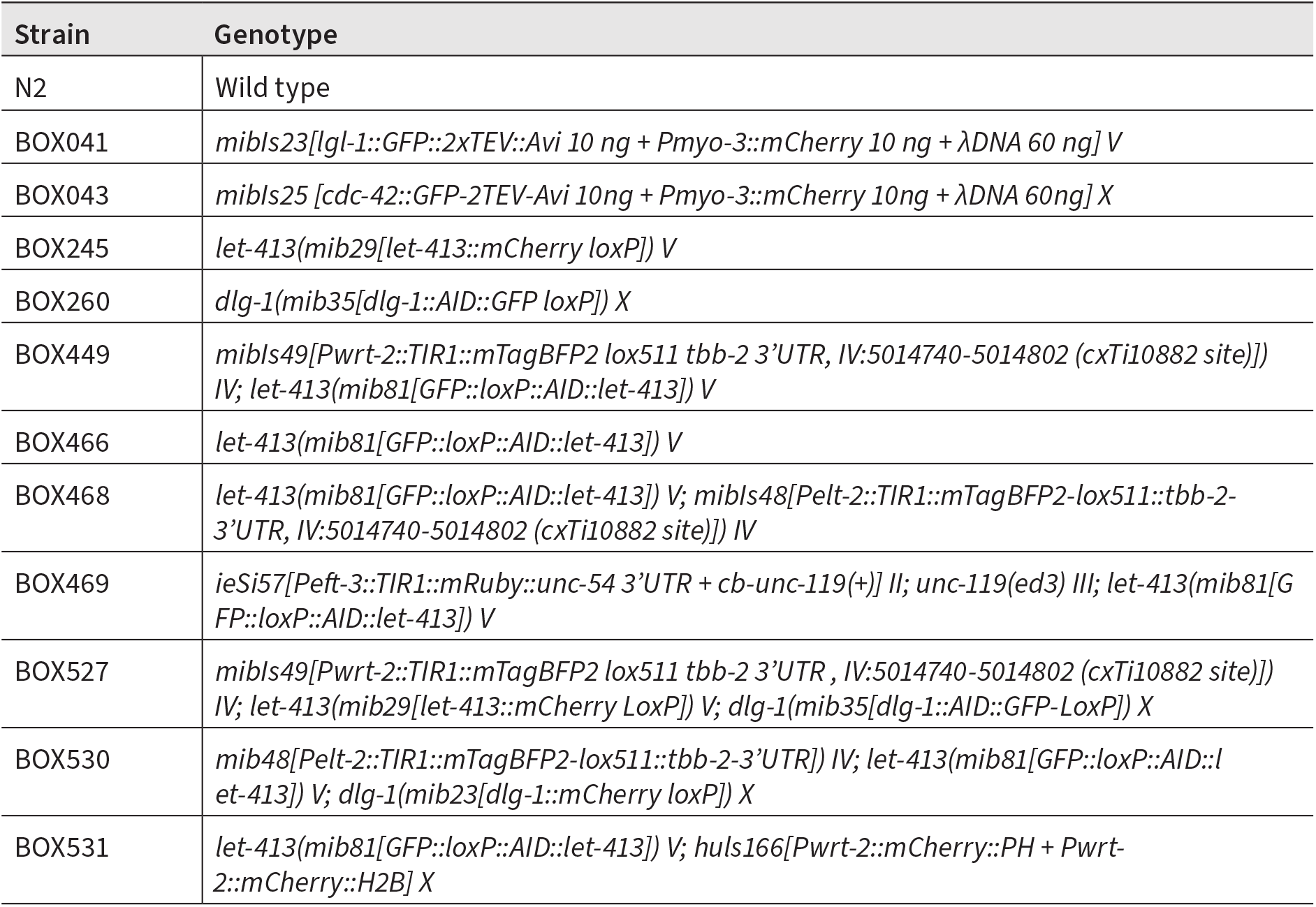
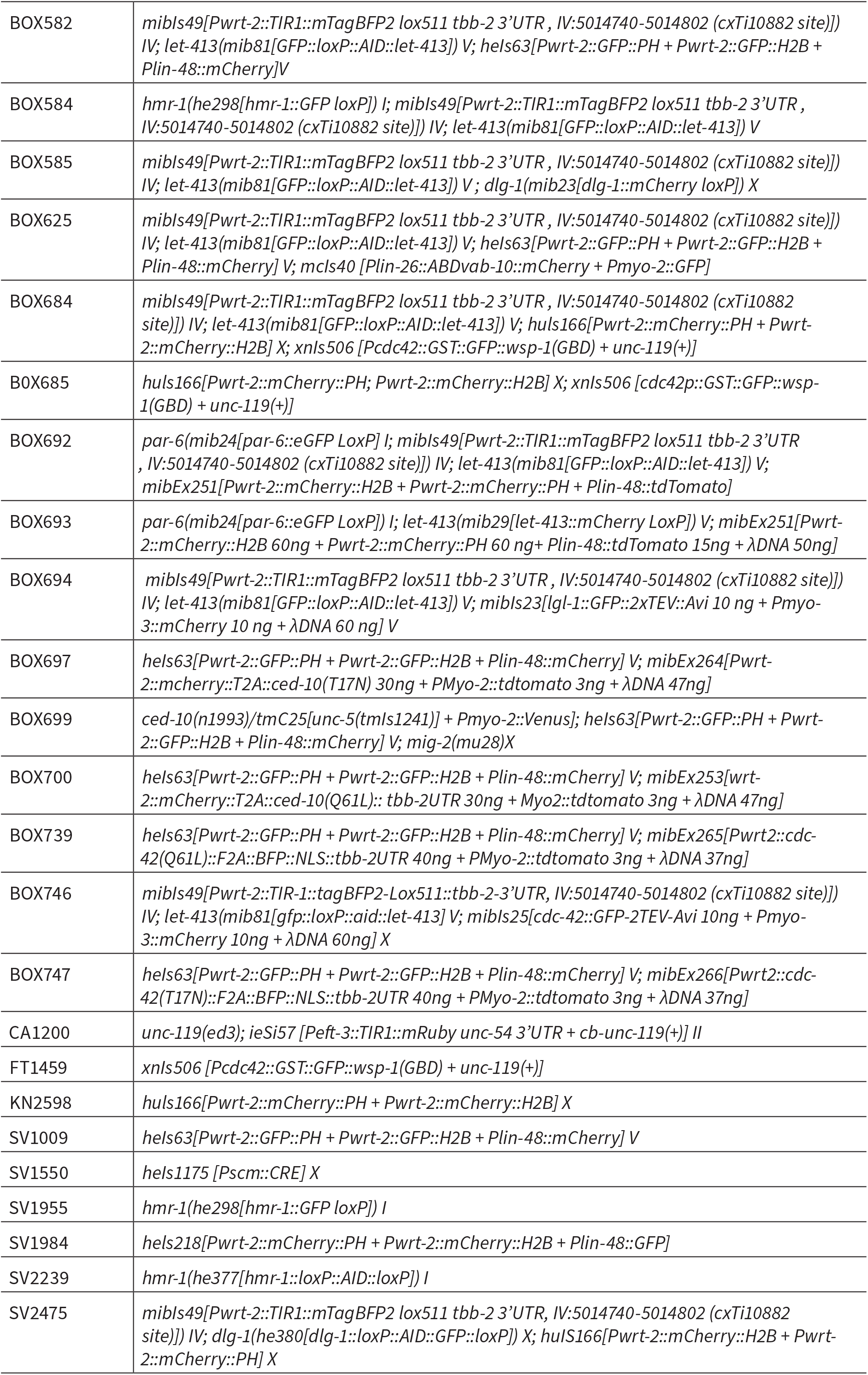
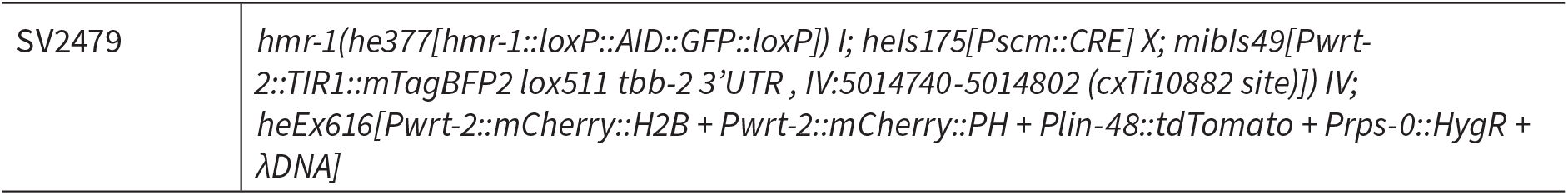
List of strains.

### *CRISPR/Cas9* genome engineering

All gene editing made use of homology-directed repair of CRISPR/Cas9-induced DNA double-strand breaks, using plasmid-based expression of Cas9 and sgRNAs. Final genomic sequences are available in Supplemental File X. All edits were made in an N2 background. Repair templates were plasmid-based, containing 190–600 bp homology arms and a self-excising cassette (SEC) for selection of candidate integrants (Dickinson et al., 2015). The *let-413::mCherry* and *dlg-1::AID::GFP* repair templates were cloned using Gibson assembly, into vector backbones pJJR83 (Addgene #75028) and pJJR82 (Addgene #75027), respectively (Gibson et al., 2009; Ramalho et al., 2020). The *GFP::AID::let-413* repair template was cloned using SapTrap assembly into vectors pDD379 (Addgene #91834) and pMLS257, respectively (Dickinson et al., 2018; Schwartz and Jorgensen, 2016). All sgRNAs were expressed from plasmids under control of a U6 promoter. To generate *GFP::AID::let-413*, the sgRNAs were incorporated into assembly vector pDD379 using SapTrap assembly. For all other sgRNAs, antisense oligonucleotide pairs were annealed and ligated into BbsI-linearized pJJR50 (Addgene #75026) (Waaijers et al., 2016). The sgRNA sequences and primers used to check integration can be found in Table 2. Injection mixes were prepared in MilliQ H_2_O and contained 50 ng/ml *Peft-3::cas9* (Addgene ID #46168) (Friedland et al., 2013) 50–100 ng/ml U6::sgRNA, and 50-75 ng/ml of repair template. All mixes also contained 2.5 ng/ml of the co-injection pharyngeal marker *Pmyo-2::GFP* or *Pmyo-2:: tdTomato* to aid in visual selection of transgenic strains. Young adult hermaphrodites were injected in the germline using an inverted micro-injection setup (Eppendorf FemtoJet 4x mounted on a Zeiss Axio Observer A.1 equipped with an Eppendorf Transferman 4r). Candidate edited progeny were selected on plates containing 250 ng/ml of hygromycin and correct genome editing was confirmed by Sanger sequencing (Macrogen Europe) of PCR amplicons encompassing the edited genomic region. From correctly edited strains, the hygromycin selection cassette was excised by a heat shock of L1 larvae at 34 °C for 1 h in a water bath.

**Table 2 –.**
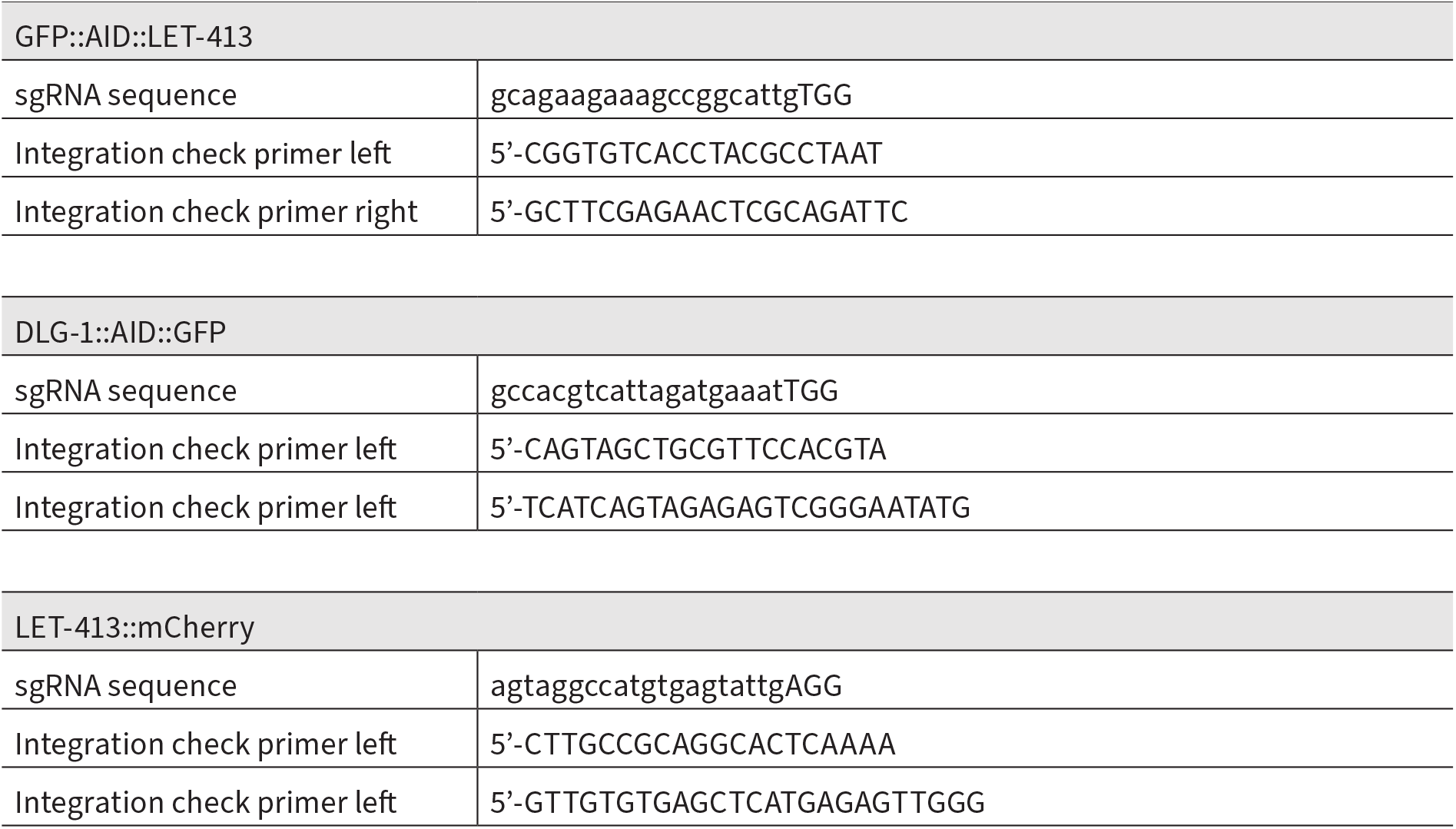
Reagents to generate CRISPR/Cas9 lines.

### CED-10 and CDC-42 Q61L and T17N expression plasmids

Plasmids for expression of constitutively active and dominant negative variants of CED-10 and CDC-42 were generated using Gibson assembly. Final plasmid sequences are available in Supplemental File 2. CED-10b and CDC-42 coding sequences carrying the appropriate base pair changes were ordered as gBlocks Gene Fragments (Integrated DNA Technologies). cdc-42 was codon optimized (Redemann et al., 2011), while ced-10 was not optimized due to inability to synthesize the gBlock for an optimized variant. Plasmids contain 2A self-cleaving peptide sequences (Szymczak and Vignali, 2005) between GTPase and fluorophore, to enable visual identification of expressing cells while minimizing the risk of altering the activity of the GTPase. Plasmids contain homology arms for insertion at the cxTi10816 Mos transposon site on chromosome IV but were only used as extrachromosomal arrays. For the generation of the lines, 20-30 ng/ul of the plasmids were injected with 50 ng/ul of lambda DNA and co-injection marker *Pmyo-2::tdTomato* (2.5 ng/ul).

### Animal synchronization and auxin treatment

NGM + auxin plates were prepared using the natural indole-3-acetic acid (IAA) from Alfa Aesar (#A10556). The stock powder was stored at 4 °C and was diluted in NGM agar cooled to 50 °C to a final concertation of 3 mM. The plates were left to dry at room temperature for 1–2 days covered with aluminum foil before seeding with OP50 bacteria. The plates were then kept at room temperature for another 1–2 days before storing at 4 °C in the dark for a maximum of 2 weeks. For auxin treatment in liquid M9, the water-soluble synthetic analog of IAA, 1-naphthaleneacetic acid (NAA), from Sigma Aldrich (#317918) was used. A 250 mM stock solution was prepared in M9 buffer and was further diluted in M9 buffer to 3 mM final concertation.

To synchronize animals, plates with eggs were washed with M9 buffer (0.22 M KH2PO4, 0.42 M Na2HPO4, 0.85 M NaCl, 0.001 M MgSO4) to remove larvae and adults but leave the eggs behind. After 1 h, plates were washed again to collect larvae hatched within that time span. Synchronized larvae were then transferred onto NMG-OP50 plates with auxin. For the depletion of DLG-1 in Fig. 5, gravid adult animals were bleached, and eggs were hatched in M9. Newly hatched larvae were kept in M9 with auxin but without food for 24 h before transferring onto NMG-OP50 plates with auxin. For non-auxin treated controls, NGM plates and M9 buffer lacking auxin were used.

### Feeding RNAi

For feeding RNAi experiments, bacteria were pre-cultured in 2 ml Lysogeny Broth (LB) supplemented with 100 µg/ml ampicillin (Amp) and 2.5 µg/ml tetracycline (Tet) at 37 °C in an incubator rotating at 200 rpm for 6-8 h, and then transferred to new tubes with a total volume of 10 ml LB for overnight culturing. To induce production of dsRNA, cultures were incubated for 90 min in the presence of 1 mM Isopropyl β-D-1-thiogalactopyranoside (IPTG). Bacterial cultures were pelleted by centrifugation at 4000 g for 15 min and resuspended in LB with 100 µg/ml ampicillin (Amp) and 2.5 µg/ml tetracycline (Tet) at 5x the original concentration. NGM agar plates supplemented with 100 µg/ml Amp and 1 mM IPTG were seeded with 250 µl of bacterial suspension, and kept at room temperature (RT) for 48 h in the dark. Six to eight L4 hermaphrodites per strain were transferred to individual NGM-RNAi plates against target genes and phenotypes were analyzed in the F1 generation.

### Larval lethality

To determine larval lethality, synchronized L1 animals were placed on NGM plates seeded with *E. coli* OP50, and either containing or lacking auxin. After 24 h animals were classified as dead or alive based on movement and response to physical touch.

### Microscopy and image processing

Imaging of *C. elegans* on agar plates for growth analysis was done using a Zeiss Axio Zoom.V16 equipped with a PlanNeoFluar Z 1x/0.25 objective and an Axiocam 506 color camera, driven by Zen Pro software. All other imaging of *C. elegans* was done by mounting embryos or larvae on a 5% agarose pad in a 10 mM Tetramisole solution in M9 buffer to induce paralysis. Nomarski DIC imaging was performed with an upright Zeiss AxioImager Z2 microscope using a 63 x 1.4 NA objective and a Zeiss AxioCam 503 monochrome camera, driven by Zeiss Zen software. Spinning disk confocal imaging was performed using a Nikon Ti-U microscope driven by MetaMorph Microscopy Automation and Image Analysis Software (Molecular Devices) and equipped with a Yokogawa CSU-X1-M1 confocal head and an Andor iXon DU-885 camera, using 60x or 100x 1.4 NA objectives. All stacks along the z-axis were obtained at 0,25 µm intervals. Maximum intensity Z projections were done in ImageJ (Fiji) software (Rueden et al., 2017; Schindelin et al., 2012). For quantifications, the same laser power and exposure times were used within experiments. Image scales were calibrated for each microscope using a micrometer slide. For display in figures, level adjustments, false coloring, and image overlays were done in Adobe Photoshop. Image rotation, cropping, and panel assembly were done in Adobe Illustrator. All edits were done non-destructively using adjustment layers and clipping masks, and images were kept in their original capture bit depth until final export from Illustrator for publication.

### Quantitative image analysis

#### *C. elegans* growth curves

Synchronized L1 animals were placed on NGM plates seeded with *E. coli* OP50 and either lacking or containing 3 mM auxin, and images were taken after 3, 7, 24, and 30 h. Animal lengths were measured by drawing a spline along the center line of the animal. All quantifications were done using ImageJ.

#### Fluorescence intensity measurements

For all fluorescence intensity measurements, mean background fluorescence levels were subtracted from measured values. Mean background intensity was determined in a circular region of ∼50 px diameter in areas within the field-of-view that did not contain any animals. Distribution plots of fluorescence intensity of GFP::AID-tagged LET-413 in the larval intestine and epidermis, as well as AID::GFP-tagged DLG-1, GFP-tagged LGL-1, and mCherry-tagged LET-413 in the epidermis were obtained by averaging the peak values of intensity profiles from 2–4 10 px-wide line-scans perpendicular to the membrane per animal. In the auxin-treated samples, where LET-413 is no longer enriched at the membrane, DLG-1::mCherry or the PH marker were used to determine the position to quantify. The intensity profiles of different animals were aligned at their peak values and trimmed manually to exclude values outside the cells/compartments of interest. For the intensity of GFP-tagged PAR-6, peak values of intensity profiles from multiple 10 px-wide line-scans in the apical side of the cytoplasm of the seam cells were obtained, averaged, and corrected for the background of the same animal. In all cases the graphs indicate the mean intensities of the values. All quantifications were done using ImageJ.

### Statistical analysis

All statistical analyses were performed using GraphPad Prism 8. For population comparisons, a D’Agostino & Pearson test of normality was first performed to determine if the data was sampled from a Gaussian distribution. For data drawn from a Gaussian distribution, comparisons between two populations were done using an unpaired t test, with Welch’s correction if the SDs of the populations differ significantly, and comparisons between >2 populations were done using a one-way ANOVA, or a Welch’s ANOVA if the SDs of the populations differ significantly. For data not drawn from a Gaussian distribution, a non-parametric test was used (Mann-Whitney for 2 populations and Kruskal-Wallis for >2 populations). ANOVA and non-parametric tests were followed up with multiple comparison tests of significance (Dunnett’s, Tukey’s, Dunnett’s T3 or Dunn’s). Tests of significance used and sample sizes are indicated in the figure legends. No statistical

method was used to pre-determine sample sizes. No samples or animals were excluded from analysis. The experiments were not randomized, and the investigators were not blinded to allocation during experiments and outcome assessment.

## Supporting information

Supplemental DNA files

## Acknowledgements and funding

We thank S. Ruijtenberg for strain SV1550, M. Soto for the *gex-2* RNAi clone, and members of the S. van den Heuvel, S. Ruijtenberg, and M. Boxem groups for helpful discussions. We also thank Wormbase (Harris et al., 2020) and the Biology Imaging Center, Faculty of Sciences, Department of Biology, Utrecht University. Some strains were provided by the Caenorhabditis Genetics Center, which is funded by NIH Office of Research Infrastructure Programs (P40 OD010440). This work was supported by the Netherlands Organization for Scientific Research (NWO)-ALW Open Program 824.14.021 and NWO-VICI 016.VICI.170.165 grants to M. Boxem, and the European Union’s Horizon 2020 research and innovation programme under the Marie Skłodowska-Curie grant agreement No. 675407 – PolarNet.

## Author Contributions

Conceptualization: A.R., J.C., S.v.d.H., M.B.; Methodology: A.R., J.C., H.R.P.; Formal analysis: A.R.; Investigation: A.R., J.C., R.S., V.G.G.; Writing - original draft: A.R., M.B.; Writing – review & editing: J.C., R.S., H.R.P., V.G.G., S.v.d.H.; Visualization: A.R., M.B.; Supervision: S.v.d.H., M.B.; Project administration: M.B.; Funding acquisition: S.v.d.H., M.B.

## References

Altun, Z. F. and Hall, D. H. (2009). Epithelial systems, hypodermis. In WormAtlas.

Audebert, S., Navarro, C., Nourry, C., Chasserot-Golaz, S., Lécine, P., Bellaiche, Y., Dupont, J.-L., Premont, R. T., Sempéré, C., Strub, J.-M., et al. (2004). Mammalian Scribble Forms a Tight Complex with the βPIX Exchange Factor. Curr. Biol. 14, 987–995.

Austin, J. and Kenyon, C. (1994). Cell contact regulates neuroblast formation in the Caenorhabditis elegans lateral epidermis. Dev. Camb. Engl. 120, 313–323.

Benton, R. and Johnston, D. S. (2003). Drosophila PAR-1 and 14-3-3 Inhibit Bazooka/PAR-3 to Establish Complementary Cortical Domains in Polarized Cells. Cell 115, 691–704.

Bilder, D., Li, M. and Perrimon, N. (2000). Cooperative regulation of cell polarity and growth by Drosophila tumor suppressors. Science 289, 113–116.

Bilder, D., Schober, M. and Perrimon, N. (2003). Integrated activity of PDZ protein complexes regulates epithelial polarity. Nat. Cell Biol. 5, 53–58.

Bone, C. R., Chang, Y.-T., Cain, N. E., Murphy, S. P. and Starr, D. A. (2016). Nuclei migrate through constricted spaces using microtubule motors and actin networks in C. elegans hypodermal cells. Dev. Camb. Engl. 143, 4193–4202.

Bonello, T. T. and Peifer, M. (2019). Scribble: A master scaffold in polarity, adhesion, synaptogenesis, and proliferation. J. Cell Biol. 218, 742–756.

Bossinger, O., Klebes, A., Segbert, C., Theres, C. and Knust, E. (2001). Zonula adherens formation in Caenorhabditis elegans requires dlg-1, the homologue of the Drosophila gene discs large. Dev Biol 230, 29–

Bossinger, O., Fukushige, T., Claeys, M., Borgonie, G. and McGhee, J. D. (2004). The apical disposition of the Caenorhabditis elegans intestinal terminal web is maintained by LET-413. Dev. Biol. 268, 448–456.

Bourne, H. R., Sanders, D. A. and McCormick, F. (1991). The GTPase superfamily: conserved structure and molecular mechanism. Nature 349, 117–127.

Brabin, C., Appleford, P. J. and Woollard, A. (2011). The Caenorhabditis elegans GATA factor ELT-1 works through the cell proliferation regulator BRO-1 and the Fusogen EFF-1 to maintain the seam stem-like fate. PLoS Genet. 7, e1002200.

Brenner, S. (1974). The genetics of Caenorhabditis elegans. Genetics 77, 71–94.

Brooks, D. R., Appleford, P. J., Murray, L. and Isaac, R. E. (2003). An essential role in molting and morphogenesis of Caenorhabditis elegans for ACN-1, a novel member of the angiotensin-converting enzyme family that lacks a metallopeptidase active site. J. Biol. Chem. 278, 52340–52346.

Cao, W., Deng, S. and Pocock, R. (2021). The UIG-1/CDC-42 guanine nucleotide exchange factor acts in parallel to CED-10/Rac1 during axon outgrowth in Caenorhabditis elegans. Small GTPases 12, 60–66.

Castiglioni, V. G., Pires, H. R., Rosas Bertolini, R., Riga, A., Kerver, J. and Boxem, M. (2020). Epidermal PAR-6 and PKC-3 are essential for larval development of C. elegans and organize non-centrosomal microtubules. eLife 9, e62067.

Chalmers, A. D., Pambos, M., Mason, J., Lang, S., Wylie, C. and Papalopulu, N. (2005). aPKC, Crumbs3 and Lgl2 control apicobasal polarity in early vertebrate development. Dev. Camb. Engl. 132, 977–986.

Chen, J., Sayadian, A.-C., Lowe, N., Lovegrove, H. E. and St Johnston, D. (2018). An alternative mode of epithelial polarity in the Drosophila midgut. PLoS Biol. 16, e3000041.

Chisholm, A. D. and Hsiao, T. I. (2012). The Caenorhabditis elegans epidermis as a model skin. I: development, patterning, and growth. Wiley Interdiscip. Rev. Dev. Biol. 1, 861–878.

Dickinson, D. J., Pani, A. M., Heppert, J. K., Higgins, C. D. and Goldstein, B. (2015). Streamlined Genome Engineering with a Self-Excising Drug Selection Cassette. Genetics 200, 1035–1049.

Dickinson, D., Slabodnick, M., Chen, A. and Goldstein, B. (2018). SapTrap assembly of repair templates for Cas9-triggered homologous recombination with a self-excising cassette. MicroPublication Biol. 2018.

Dow, L. E., Brumby, A. M., Muratore, R., Coombe, M. L., Sedelies, K. A., Trapani, J. A., Russell, S. M., Richardson, H. E. and Humbert, P. O. (2003). hScrib is a functional homologue of the Drosophila tumour suppressor Scribble. Oncogene 22, 9225–9230.

Dow, L. E., Kauffman, J. S., Caddy, J., Peterson, A. S., Jane, S. M., Russell, S. M. and Humbert, P. O. (2007). The tumour-suppressor Scribble dictates cell polarity during directed epithelial migration: regulation of Rho GTPase recruitment to the leading edge. Oncogene 26, 2272–2282.

Dyer, J. O., Demarco, R. S. and Lundquist, E. A. (2010). Distinct roles of Rac GTPases and the UNC-73/Trio and PIX-1 Rac GTP exchange factors in neuroblast protrusion and migration in C. elegans. Small GTPases 1, 44–61.

Elsum, I., Yates, L., Humbert, P. O. and Richardson, H. E. (2012). The Scribble-Dlg-Lgl polarity module in development and cancer: from flies to man. Essays Biochem. 53, 141–168.

Friedland, A. E., Tzur, Y. B., Esvelt, K. M., Colaiácovo, M. P., Church, G. M. and Calarco, J. A. (2013). Heritable genome editing in C. elegans via a CRISPR-Cas9 system. Nat. Methods 10, 741–743.

Gally, C., Wissler, F., Zahreddine, H., Quintin, S., Landmann, F. and Labouesse, M. (2009). Myosin II regulation during C. elegans embryonic elongation: LET-502/ROCK, MRCK-1 and PAK-1, three kinases with different roles. Dev. Camb. Engl. 136, 3109–3119.

Gateff, E. and Schneiderman, H. A. (1974). Developmental capacities of benign and malignant neoplasms of Drosophila. Wilhelm Roux Arch. Entwicklungsmechanik Org. 176, 23–65.

Gibson, D. G., Young, L., Chuang, R.-Y., Venter, J. C., Hutchison, C. A. and Smith, H. O. (2009). Enzymatic assembly of DNA molecules up to several hundred kilobases. Nat. Methods 6, 343–345.

Grifoni, D., Garoia, F., Bellosta, P., Parisi, F., De Biase, D., Collina, G., Strand, D., Cavicchi, S. and Pession, A. (2007). aPKCzeta cortical loading is associated with Lgl cytoplasmic release and tumor growth in Drosophila and human epithelia. Oncogene 26, 5960–5965.

Khoury, M. J. and Bilder, D. (2020). Distinct activities of Scrib module proteins organize epithelial polarity. Proc. Natl. Acad. Sci. U. S. A. 117, 11531–11540.

Köppen, M., Simske, J. S., Sims, P. A., Firestein, B. L., Hall, D. H., Radice, A. D., Rongo, C. and Hardin, J. D. (2001). Cooperative regulation of AJM-1 controls junctional integrity in Caenorhabditis elegans epithelia. Nat. Cell Biol. 3, 983–991.

Kumfer, K. T., Cook, S. J., Squirrell, J. M., Eliceiri, K. W., Peel, N., O’Connell, K. F. and White, J. G. (2010). CGEF-1 and CHIN-1 regulate CDC-42 activity during asymmetric division in the Caenorhabditis elegans embryo. Mol. Biol. Cell 21, 266–277.

Laprise, P., Beronja, S., Silva-Gagliardi, N. F., Pellikka, M., Jensen, A. M., McGlade, C. J. and Tepass, U. (2006). The FERM Protein Yurt Is a Negative Regulatory Component of the Crumbs Complex that Controls Epithelial Polarity and Apical Membrane Size. Dev. Cell 11, 363–374.

Legouis, R., Gansmuller, A., Sookhareea, S., Bosher, J. M., Baillie, D. L. and Labouesse, M. (2000). LET-413 is a basolateral protein required for the assembly of adherens junctions in Caenorhabditis elegans. Nat. Cell Biol. 2, 415.

Legouis, R., Jaulin-Bastard, F., Schott, S., Navarro, C., Borg, J.-P. and Labouesse, M. (2003). Basolateral targeting by leucine-rich repeat domains in epithelial cells. EMBO Rep. 4, 1096–1102.

Liu, H., Wang, S., Hang, W., Gao, J., Zhang, W., Cheng, Z., Yang, C., He, J., Zhou, J., Chen, J., et al. (2018). LET-413/Erbin acts as a RAB-5 effector to promote RAB-10 activation during endocytic recycling. J. Cell Biol. 217, 299–314.

Lundquist, E. A., Reddien, P. W., Hartwieg, E., Horvitz, H. R. and Bargmann, C. I. (2001). Three C. elegans Rac proteins and several alternative Rac regulators control axon guidance, cell migration and apoptotic cell phagocytosis. Dev. Camb. Engl. 128, 4475–4488.

McMahon, L., Legouis, R., Vonesch, J.-L. and Labouesse, M. (2001). Assembly of C. elegans apical junctions involves positioning and compaction by LET-413 and protein aggregation by the MAGUK protein DLG-1. J. Cell Sci. 114, 2265–2277.

Mechler, B. M., McGinnis, W. and Gehring, W. J. (1985). Molecular cloning of lethal(2)giant larvae, a recessive oncogene of Drosophila melanogaster. EMBO J. 4, 1551–1557.

Michaelis, U. R., Chavakis, E., Kruse, C., Jungblut, B., Kaluza, D., Wandzioch, K., Manavski, Y., Heide, H., Santoni, M.-J., Potente, M., et al. (2013). The polarity protein Scrib is essential for directed endothelial cell migration. Circ. Res. 112, 924–934.

Mullaney, B. C., Blind, R. D., Lemieux, G. A., Perez, C. L., Elle, I. C., Faergeman, N. J., Van Gilst, M. R., Ingraham, H. A. and Ashrafi, K. (2010). Regulation of C. elegans fat uptake and storage by acyl-CoA synthase-3 is dependent on NR5A family nuclear hormone receptor nhr-25. Cell Metab. 12, 398–410.

Nishimura, K., Fukagawa, T., Takisawa, H., Kakimoto, T. and Kanemaki, M. (2009). An auxin-based degron system for the rapid depletion of proteins in nonplant cells. Nat. Methods 6, 917–922.

Nola, S., Sebbagh, M., Marchetto, S., Osmani, N., Nourry, C., Audebert, S., Navarro, C., Rachel, R., Montcouquiol, M., Sans, N., et al. (2008). Scrib regulates PAK activity during the cell migration process. Hum. Mol. Genet. 17, 3552–3565.

Osmani, N., Vitale, N., Borg, J.-P. and Etienne-Manneville, S. (2006). Scrib Controls Cdc42 Localization and Activity to Promote Cell Polarization during Astrocyte Migration. Curr. Biol. 16, 2395–2405.

Osmani, N., Peglion, F., Chavrier, P. and Etienne-Manneville, S. (2010). Cdc42 localization and cell polarity depend on membrane traffic. J. Cell Biol. 191, 1261–1269.

Ouellette, M.-H., Martin, E., Lacoste-Caron, G., Hamiche, K. and Jenna, S. (2016). Spatial control of active CDC-42 during collective migration of hypodermal cells in Caenorhabditis elegans. J. Mol. Cell Biol. 8, 313– 327.

Pannekoek, W.-J., de Rooij, J. and Gloerich, M. (2019). Force transduction by cadherin adhesions in morphogenesis. F1000Research 8.

Patel, F. B., Bernadskaya, Y. Y., Chen, E., Jobanputra, A., Pooladi, Z., Freeman, K. L., Gally, C., Mohler, W. A. and Soto, M. C. (2008). The WAVE/SCAR complex promotes polarized cell movements and actin enrichment in epithelia during C. elegans embryogenesis. Dev. Biol. 324, 297–309.

Peters, E. C., Gossett, A. J., Goldstein, B., Der, C. J. and Reiner, D. J. (2013). Redundant canonical and noncanonical Caenorhabditis elegans p21-activated kinase signaling governs distal tip cell migrations. G3 Bethesda Md 3, 181–195.

Pilipiuk, J., Lefebvre, C., Wiesenfahrt, T., Legouis, R. and Bossinger, O. (2009). Increased IP3/Ca2+ signaling compensates depletion of LET-413/DLG-1 in C. elegans epithelial junction assembly. Dev. Biol. 327, 34–47.

Pinheiro, D. and Bellaïche, Y. (2018). Mechanical Force-Driven Adherens Junction Remodeling and Epithelial Dynamics. Dev. Cell 47, 3–19.

Qin, Y., Capaldo, C., Gumbiner, B. M. and Macara, I. G. (2005). The mammalian Scribble polarity protein regulates epithelial cell adhesion and migration through E-cadherin. J. Cell Biol. 171, 1061–1071.

Ramalho, J. J., Sepers, J. J., Nicolle, O., Schmidt, R., Cravo, J., Michaux, G. and Boxem, M. (2020). C-terminal phosphorylation modulates ERM-1 localization and dynamics to control cortical actin organization and support lumen formation during Caenorhabditis elegans development. Development 147, dev188011.

Raman, R., Damle, I., Rote, R., Banerjee, S., Dingare, C. and Sonawane, M. (2016). aPKC regulates apical localization of Lgl to restrict elongation of microridges in developing zebrafish epidermis. Nat. Commun. 7, 11643.

Rauzi, M. (2020). Cell intercalation in a simple epithelium. Philos. Trans. R. Soc. Lond. B. Biol. Sci. 375, 20190552.

Reddien, P. W. and Horvitz, H. R. (2000). CED-2/CrkII and CED-10/Rac control phagocytosis and cell migration in Caenorhabditis elegans. Nat. Cell Biol. 2, 131–136.

Redemann, S., Schloissnig, S., Ernst, S., Pozniakowsky, A., Ayloo, S., Hyman, A. A. and Bringmann, H. (2011). Codon adaptation–based control of protein expression in C. elegans. Nat. Methods 8, 250–252.

Rella, L., Fernandes Póvoa, E. E. and Korswagen, H. C. (2016). The Caenorhabditis elegans Q neuroblasts: A powerful system to study cell migration at single-cell resolution in vivo. Genes. N. Y. N 2000 54, 198–211.

Ridley, A. J. (2011). Life at the leading edge. Cell 145, 1012–1022.

Rueden, C. T., Schindelin, J., Hiner, M. C., DeZonia, B. E., Walter, A. E., Arena, E. T. and Eliceiri, K. W. (2017). ImageJ2: ImageJ for the next generation of scientific image data. BMC Bioinformatics 18, 529.

Santoni, M.-J., Kashyap, R., Camoin, L. and Borg, J.-P. (2020). The Scribble family in cancer: twentieth anniversary. Oncogene 39, 7019–7033.

Sasidharan, S., Borinskaya, S., Patel, F., Bernadskaya, Y., Mandalapu, S., Agapito, M. and Soto, M. C. (2018). WAVE regulates Cadherin junction assembly and turnover during epithelial polarization. Dev. Biol. 434, 133–148.

Schaks, M., Giannone, G. and Rottner, K. (2019). Actin dynamics in cell migration. Essays Biochem. 63, 483– 495.

Schindelin, J., Arganda-Carreras, I., Frise, E., Kaynig, V., Longair, M., Pietzsch, T., Preibisch, S., Rueden, C., Saalfeld, S., Schmid, B., et al. (2012). Fiji: an open-source platform for biological-image analysis. Nat. Methods 9, 676–682.

Schwartz, M. L. and Jorgensen, E. M. (2016). SapTrap, a Toolkit for High-Throughput CRISPR/Cas9 Gene Modification in Caenorhabditis elegans. Genetics 202, 1277–1288.

Segbert, C., Johnson, K., Theres, C., van Furden, D. and Bossinger, O. (2004). Molecular and functional analysis of apical junction formation in the gut epithelium of Caenorhabditis elegans. Dev Biol 266, 17–26.

Shakir, M. A., Gill, J. S. and Lundquist, E. A. (2006). Interactions of UNC-34 Enabled with Rac GTPases and the NIK kinase MIG-15 in Caenorhabditis elegans axon pathfinding and neuronal migration. Genetics 172, 893–913.

Shao, J., He, K., Wang, H., Ho, W. S., Ren, X., An, X., Wong, M. K., Yan, B., Xie, D., Stamatoyannopoulos, J., et al. (2013). Collaborative regulation of development but independent control of metabolism by two epidermis-specific transcription factors in Caenorhabditis elegans. J. Biol. Chem. 288, 33411–33426.

Silhánková, M., Jindra, M. and Asahina, M. (2005). Nuclear receptor NHR-25 is required for cell-shape dynamics during epidermal differentiation in Caenorhabditis elegans. J. Cell Sci. 118, 223–232.

Soto, M. C., Qadota, H., Kasuya, K., Inoue, M., Tsuboi, D., Mello, C. C. and Kaibuchi, K. (2002). The GEX-2 and GEX-3 proteins are required for tissue morphogenesis and cell migrations in C. elegans. Genes Dev. 16, 620–632.

Stephens, R., Lim, K., Portela, M., Kvansakul, M., Humbert, P. O. and Richardson, H. E. (2018). The Scribble Cell Polarity Module in the Regulation of Cell Signaling in Tissue Development and Tumorigenesis. J. Mol. Biol. 430, 3585–3612.

Sulston, J. E. and Horvitz, H. R. (1977). Post-embryonic cell lineages of the nematode, Caenorhabditis elegans. Dev. Biol. 56, 110–156.

Sun, T., Yang, L., Kaur, H., Pestel, J., Looso, M., Nolte, H., Krasel, C., Heil, D., Krishnan, R. K., Santoni, M.-J., et al. (2017). A reverse signaling pathway downstream of Sema4A controls cell migration via Scrib. J. Cell Biol. 216, 199–215.

Szymczak, A. L. and Vignali, D. A. A. (2005). Development of 2A peptide-based strategies in the design of multicistronic vectors. Expert Opin. Biol. Ther. 5, 627–638.

Tanentzapf, G. and Tepass, U. (2003). Interactions between the crumbs, lethal giant larvae and bazooka pathways in epithelial polarization. Nat. Cell Biol. 5, 46.

van der Horst, S. E. M., Cravo, J., Woollard, A., Teapal, J. and van den Heuvel, S. (2019). C. elegans Runx/ CBFβ suppresses POP-1 TCF to convert asymmetric to proliferative division of stem cell-like seam cells. Dev. Camb. Engl. 146.

Waaijers, S., Muñoz, J., Berends, C., Ramalho, J. J., Goerdayal, S. S., Low, T. Y., Zoumaro-Djayoon, A. D., Hoffmann, M., Koorman, T., Tas, R. P., et al. (2016). A tissue-specific protein purification approach in Caenorhabditis elegans identifies novel interaction partners of DLG-1/Discs large. BMC Biol. 14, 66.

Walck-Shannon, E. and Hardin, J. (2014). Cell intercalation from top to bottom. Nat. Rev. Mol. Cell Biol. 15, 34–48.

Walck-Shannon, E., Reiner, D. and Hardin, J. (2015). Polarized Rac-dependent protrusions drive epithelial intercalation in the embryonic epidermis of C. elegans. Dev. Camb. Engl. 142, 3549–3560.

Walck-Shannon, E., Lucas, B., Chin-Sang, I., Reiner, D., Kumfer, K., Cochran, H., Bothfeld, W. and Hardin, J. (2016). CDC-42 Orients Cell Migration during Epithelial Intercalation in the Caenorhabditis elegans Epidermis. PLoS Genet. 12, e1006415.

Wallace, A. G., Raduwan, H., Carlet, J. and Soto, M. C. (2018). The RhoGAP HUM-7/Myo9 integrates signals to modulate RHO-1/RhoA during embryonic morphogenesis in Caenorhabditiselegans. Dev. Camb. Engl. 145.

Ward, J. D., Bojanala, N., Bernal, T., Ashrafi, K., Asahina, M. and Yamamoto, K. R. (2013). Sumoylated NHR-25/NR5A regulates cell fate during C. elegans vulval development. PLoS Genet. 9, e1003992.

Warner, H., Wilson, B. J. and Caswell, P. T. (2019). Control of adhesion and protrusion in cell migration by Rho GTPases. Curr. Opin. Cell Biol. 56, 64–70.

Welchman, D. P., Mathies, L. D. and Ahringer, J. (2007). Similar requirements for CDC-42 and the PAR-3/ PAR-6/PKC-3 complex in diverse cell types. Dev. Biol. 305, 347–357.

Wildwater, M., Sander, N., de Vreede, G. and van den Heuvel, S. (2011). Cell shape and Wnt signaling redundantly control the division axis of C. elegans epithelial stem cells. Development 138, 4375–4385.

Woods, D. F. and Bryant, P. J. (1989). Molecular cloning of the lethal(1)discs large-1 oncogene of Drosophila. Dev. Biol. 134, 222–235.

Yamanaka, T., Horikoshi, Y., Izumi, N., Suzuki, A., Mizuno, K. and Ohno, S. (2006). Lgl mediates apical domain disassembly by suppressing the PAR-3-aPKC-PAR-6 complex to orient apical membrane polarity. J. Cell Sci. 119, 2107–2118.

Zhang, L., Ward, J. D., Cheng, Z. and Dernburg, A. F. (2015). The auxin-inducible degradation (AID) system enables versatile conditional protein depletion in C. elegans. Dev. Camb. Engl. 142, 4374–4384.

Zilberman, Y., Abrams, J., Anderson, D. C. and Nance, J. (2017). Cdc42 regulates junctional actin but not cell polarization in the Caenorhabditis elegans epidermis. J. Cell Biol.

