## Supplemental Figures for "*Caenorhabditis elegans* LET-413 Scribble is essential in the epidermis for growth, viability, and directional outgrowth of epithelial seam cells"

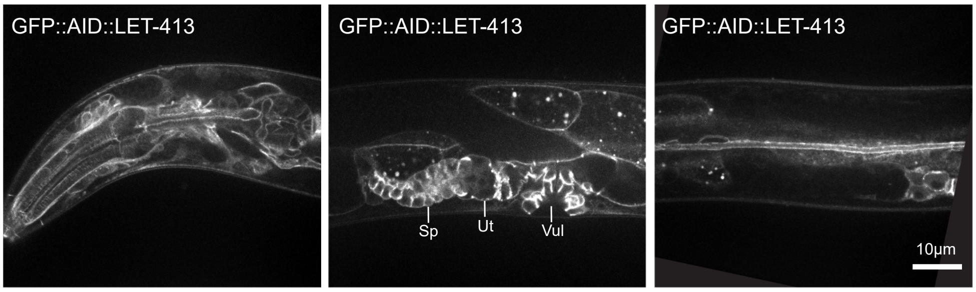


**Figure S1. Larval expression of LET-413.** Expression of GFP::AID::LET-413 in the pharynx (left), reproductive system (middle), and excretory canal (right) of *GFP::AID::let-413* animals. Sp: spermatheca, Ut: Uterus, Vul: Vulva. Related to Figure 1.

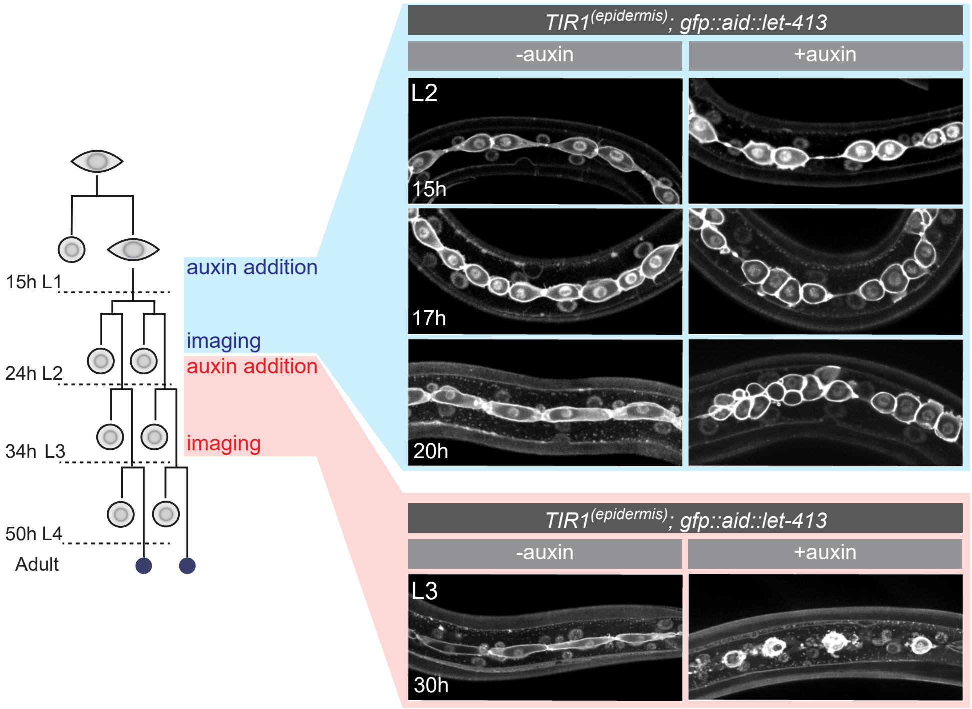


**Figure S2. LET-413 is required for seam cell outgrowth throughout development.** Time series of L2 and L3 seam cells divisions and subsequent extension in LET-413-depleted (+auxin) or control animals (-auxin). Seam-specific GFP::H2B and GFP::PH mark DNA and cell membrane, respectively. For the L2 division pattern (blue), auxin was added from 8h after hatching, and for the L3 division pattern (red) from 19h after hatching. Related to Figure 3.

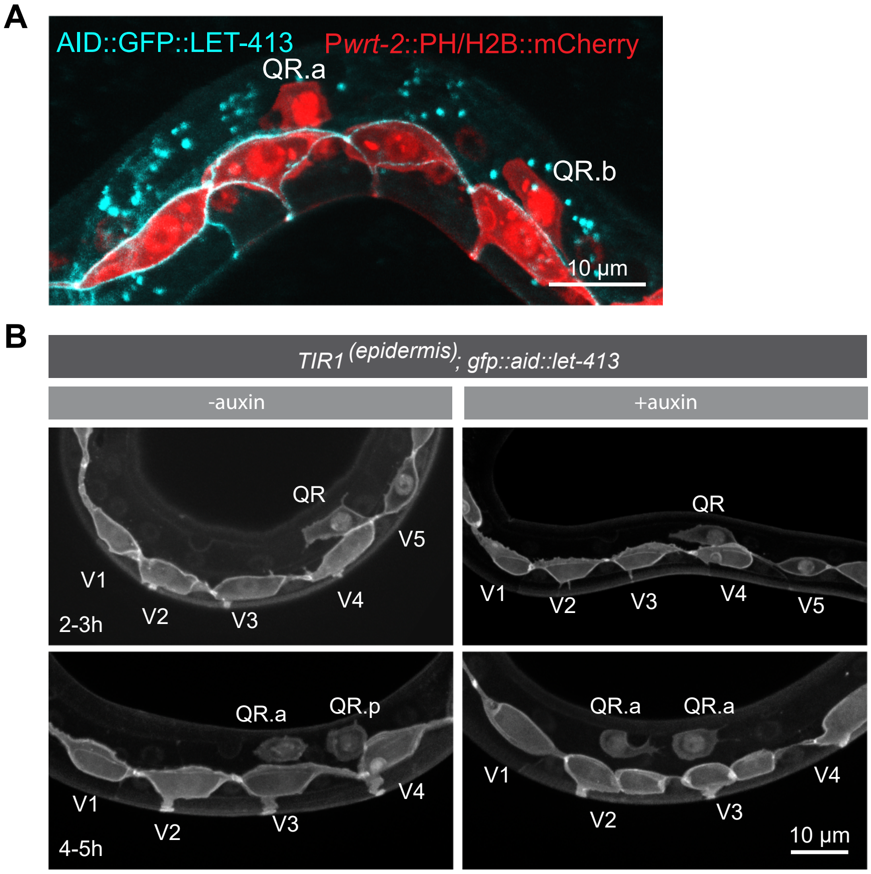


**Figure S3**. **Degradation of LET-413 in the epidermis does not affect Q cell migration. (A)** Q cell descendant QRa and QRb during anterior migration, marked with epidermal-specific mCherry::H2B and mCherry::PH. No expression of GFP::AID::LET-413 is detected. **(B)** Migration and division of Q cell descendants in LET-413-depleted (+auxin) or control animals (-auxin) at 2–3 h and 4–5 h post hatching. Related to Figure 3.

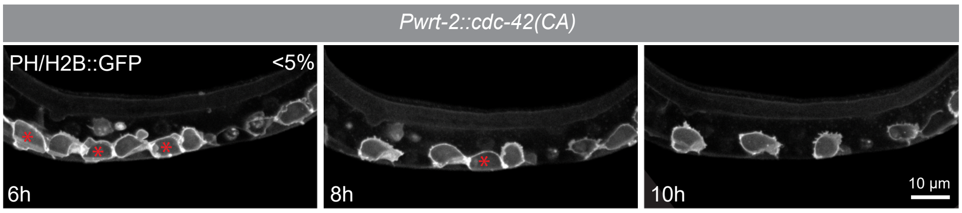


**Figure S4**. Time series of the distribution of the PH::GFP and H2B::GFP marker in animals expressing a constitutively active (CA) form of CDC-42 at 6, 8 and 10 h post hatching. ~5% of the imaged animals show this phenotype in the L1. Asterisks indicate cells that will fuse with the hypodermis. Related to Figure 6.

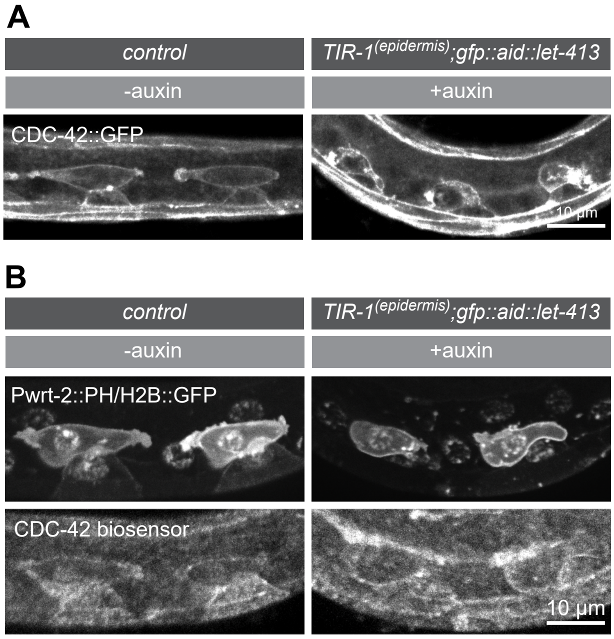


**Figure S5**. (**A**) Distribution of CDC-42::GFP marker in control and LET-413-depleted animals at 8 h post hatching. Genotypes are *cdc-42::GFP* for the control and *Pwrt-2::TIR1::BFP; GFP::AID::let-413; cdc-42::GFP* for the auxin treated animals. (**B**) Distribution of PH::GFP and H2B::GFP markers and the CDC-42 biosensor in control and LET-413-depleted animals at 8 h post hatching. Genotypes are *Pwrt2::PH::mCherry Pwrt-2::H2B::mCherry; Pcdc-42::GST::GFP::wsp-1(GBD)* for the control and *Pwrt-2::TIR1::BFP; GFP::AID::let-413; Pwrt2::PH::mCherry Pwrt-2::H2B::mCherry; Pcdc-42::GST::GFP::wsp-1(GBD)* for the auxin-treated animals. Related to Figure 6.
